# Stress-adapted codon usage enables efficient virulence-gene expression in *Salmonella*

**DOI:** 10.64898/2026.02.22.707322

**Authors:** Frédéric Goormaghtigh, Dirk Bumann

## Abstract

Virulence genes in bacterial pathogens are often A/T-rich and horizontally acquired, yet they must be translated efficiently under stress and nutrient limitation encountered during infection. Here we show that synonymous codon usage contributes directly to this problem. Using CodonPipe, a genome-scale framework for synonymous codon-usage analysis, we find that bacterial genomes contain functionally organized codon-usage landscapes that extend well beyond previous knowledge. In *Salmonella enterica*, virulence genes form a codon signature that is distinct from ribosomal genes and other mobile elements. This signature favors wobble decoding, increased use of rare tRNA isoacceptors, and enrichment of codons previously shown to better preserve translation during amino-acid limitation. Using codon-recoded fluorescent reporters, we show that virulence codon usage outperforms ribosomal codon usage selectively during nutrient starvation and, more strongly, in host-mimicking conditions. In mouse infections, overexpression of virulence-recoded reporters imposes a strong selective cost. Comparative analyses further indicate that related decoding signatures are conserved across diverse Enterobacteriaceae pathogens. These findings support stress-adapted codon usage as a mechanism that promotes virulence-gene expression during infection. More broadly, our work contributes to explain the long-observed A/T-rich codon bias of virulence genes and identifies tRNA charging and modifications as promising broad anti-virulence targets.

## Introduction

Eighteen of the twenty amino acids incorporated during translation are encoded by multiple synonymous codons, yet these codons are not used equally (1). Across organisms, codon usage varies with nucleotide composition, gene expression level, tRNA availability and evolutionary history, thereby shaping how genes are translated and how genomes evolve. In bacteria, this variation has classically been interpreted through three broad signatures: codon optimized highly expressed genes, codon biased horizontally acquired genes and the more heterogeneous core genome.

Highly expressed genes, including ribosomal proteins and other components of the translational machinery preferentially use codons matching abundant tRNAs. This is thought to improve translational efficiency and accuracy while limiting ribosome stalling and depletion of charged tRNAs (1). This principle also underlies classical metrics such as the Codon Adaptation Index, which uses highly expressed genes as a reference to estimate translational adaptation and predict gene expression levels (2). In contrast, horizontally acquired genes often display atypical, A/T-rich codon usage. This may reflect donor-genome composition, more efficient transfer of AT-rich genes, or preferential retention of acquired genes that can be silenced outside of host environments by the AT-binding protein H-NS (3–5).

This classical framework is powerful but incomplete as evidenced by the fact that codon usage does not necessarily correlate with gene expression (6–8). Synonymous codon changes can alter mRNA folding near the ribosome-binding site and subsequently transcript stability (6–8). Importantly, translation initiation rather than elongation is often rate-limiting, and many genes encode an N-terminal “ramp” of slowly translated codons that regulates ribosomal trafficking (9,10). Moreover, codon effects can become most pronounced under specific physiological conditions, when amino acid availability, tRNA charging or translational demand change (11–13). During amino acid limitation, tRNA isoacceptors carrying the same amino acid can differ substantially in charging state, generating codons that are comparatively sensitive or robust to starvation (11,12). Thus, codon usage may reflect not only a static signature of gene origin or expression level but may also encode adaptation to specific physiological conditions. This might be especially relevant for bacterial pathogens. Many virulence genes are horizontally acquired, A/T-rich and strongly biased in synonymous codon usage. Yet they are among the most highly expressed genes during infection (14), when bacteria face host-imposed stresses such as nutrient deprivation, acidic pH and immune attacks.

Here, we asked whether virulence-gene codon bias contributes functionally to expression under host-imposed stress, rather than merely reflecting horizontal acquisition. We therefore developed CodonPipe, a genome-wide codon-usage analysis framework that represents genes in synonymous codon-usage space, identifies functional clusters and relates codon signatures to predicted decoding strategies. This revealed that codon usage is far more complex and functionally diverse than the three classical signatures. Across bacteria and yeast, genes sharing cellular functions converged toward specific codon-usage signatures, including ribosomal proteins, aminoacyl-tRNA synthetases, carbon metabolism genes, mobile genetic elements, virulence genes and many others. This suggests that synonymous codon choice can encode functional and physiological information beyond expression level, nucleotide composition or gene origin.

Virulence genes provided a particularly compelling example of this principle. In *Salmonella enterica*, they formed a coherent A/T-rich codon signature that was distinct from ribosomal genes and from other horizontally acquired elements. This signature favored wobble decoding and rare tRNA isoacceptor usage, and was enriched in codons predicted from prior studies to better preserve translation during amino-acid limitation (11,12). Using recoded fluorescent reporters and mouse co-infection studies, we provide evidence that virulence-like codon usage enhances expression under infection-relevant nutrient stress, while minimizing impact on translation homeostasis. Comparative analyses further showed that virulence genes from diverse Enterobacteriaceae pathogens converge toward related codon-usage and decoding signatures despite differences in function, genomic location and evolutionary origin, suggesting that this strategy may represent a broader principle of bacterial pathogenesis. Together, these findings support a model in which virulence codon bias can function as a stress-adapted decoding strategy, while remaining compatible with established roles for horizontal acquisition and H-NS-mediated silencing (3,5). Our findings also identify tRNA charging and tRNA modifications as promising targets for broad anti-virulence strategies.

## Materials and Methods

### Microbiology and Molecular Biology

Bacteria were grown in Lennox lysogeny broth (LB), M9 or MES-ch (SPI-2 inducing medium). M9 medium contained 1× M9 salts, 0.4% glucose and 2 mM MgSO4, and was supplemented with 0.2% casamino acids where indicated. MES-ch was prepared as previously described (15) and contained 100 mM MES-KOH pH 5.5, 5 mM KCl, 15 mM NH_4_Cl, 0.5 mM K_2_SO_4_, 0.113 mM KH_2_PO_4_, 0.05 mM MgSO_4_, 0.01% glucose, 0.02% glycerol, supplemented with casamino acids 0.2% where indicated. Preparation of MES-ch is described in Table S3. All media contained kanamycin (50 µg/mL) to maintain plasmid selection.

All strains were derived from SME51 (16), a prototrophic hisG^Leu69^ derivative of *Salmonella enterica* serovar Typhimurium SL1344 (further referred to as *STm*) (17). Strains and plasmids used in this study are described in Table S4. Vir- and HEG-recoded fluorescent protein coding sequences are available at Data S2. Full plasmid DNA sequences are available at Data S2. Fluorescent reporter plasmids are based on low-copy pSC101 derivatives carrying a kanamycin resistance cassette.

- FreGo911-914 plasmids carry (i) vir- or HEG-recoded *gfpmut2* under the control of the proD constitutive promoter (18) and (ii) a codon-optimized (using IDT online tool) *mScarlet-i3* gene under the control of the constitutive PybaJ promoter (14), serving as an internal control.
- FreGo1200 and FreGo2000 plasmids comprise (i) a SPI-2 (*PssaG* - FreGo1200) or arabinose (pBAD – FreGo2000) inducible promoter controlling expression of vir- or HEG-recoded fluorescent protein genes *mCherry*, *mtagBFP2* or *ypet* together with (ii) the constitutive promoter PybaJ (14) controlling expression of a codon-optimized (using IDT online tool) fluorescent protein gene (*mCherry*, *mtagBFP2*, *mNeonGreen* or *ypet*), serving as an internal fluorescent control and used to discriminate strains expressing recoded fluorescent proteins during mice experiments. Recoded fluorescent protein genes were cloned with an -LVA C-terminal degradation tag (19) to avoid possible bias by sequestering large amounts of free amino acids in stable fluorescent proteins during induction of recoded fluorescent protein genes.

### Codon engineering of fluorescent protein genes

Fluorescent protein (FP) genes were recoded to match codon usage of either virulence genes (vir) or highly expressed ribosomal genes (HEG) in *STm*. Codon usage tables were generated for virulence and ribosomal gene clusters and used as recoding references. Because codon identity within the N-terminal coding region can affect translational ramp formation, mRNA stability, and initiation efficiency (6,9,10,20), the first 25 codons were optimized using the IDT codon-usage tool and kept identical between paired vir- and HEG-recoded variants. The remainder of each coding sequence was recoded using a MATLAB-based pipeline available at Zenodo (https://doi.org/10.5281/zenodo.17072593). This pipeline takes a coding sequence and a target codon-usage table as inputs and generates a recoded sequence while preserving amino acid identity. It also calculates distance metrics to the original and target codon biases. Recoded fluorescent protein genes coding sequences are available in Data S2.

### Comparison of recoded fluorescent proteins

To trigger casamino acid starvation, bacteria were grown overnight in M9 or MES-ch medium supplemented with 0.2% casamino acids and kanamycin (50 µg/mL). The following morning, cultures were diluted 1:100 into the same fresh medium and grown for 4 h to mid-exponential phase. Bacteria were pelleted and washed twice in PBS to minimize casamino-acid carryover. Bacteria were then diluted to a final OD_600_ of 0.02 in a 24-well microplate prefilled with 1 mL of M9 or MES-ch medium either lacking casamino acids (starvation condition) or supplemented with 0.2% casamino acids (non-starvation control). Plates were incubated at 37°C with continuous shaking in a Tecan Spark or Spectramax i5 plate reader, with OD_600_ and fluorescence recorded every 15 min over 18 h. Fluorescence was measured using the appropriate excitation and emission filters: *mtagBFP2* Ex399/20 Em454/20; *gfpmut2* and *ypet* Ex500/20 Em545/20; *mScarlet-i3* and *mCherry* Ex570/20 Em620/20. Absorbance was measured at 600 nm. Full datasets for all microplate reader experiments are provided in Data S6.

Fluorescence data were analyzed by first normalizing fluorescence intensity to OD_600_. Then, the vir/HEG fluorescence ratio was calculated and plotted over time. For FreGo911-914 plasmids, which constitutively express recoded *gfpmut2* variants, vir/HEG ratios reproducibly peaked between 3 and 6 h. By contrast, FreGo2000 plasmids, which drive arabinose-inducible expression of recoded fluorescent proteins, peaked between 1 and 3h, consistent with de novo induction of fluorescent proteins synthesis rather than replenishment of proteins that had accumulated before transfer to starvation medium. In in vivo mimicking and SPI-2 inducing MES-ch medium, these peaks shifted to later time points, consistent with slower growth (Figure S9) and reduced availability of essential nutrients (phosphate, magnesium, carbon sources), which likely affect translation efficiency. For statistical comparisons and for bar graphs in Figure 5c, average vir/HEG fluorescence ratios were determined over the indicated time windows, providing robust metrics for comparison between independent biological replicates. Finally, to generate time profiles shown in Figure 5d, vir/HEG fluorescence ratios measured during starvation were normalized to the corresponding vir/HEG ratios in MES-ch medium supplemented with 0.2% casamino acids.

Relative fitness in vitro (Figure 5f) was determined as the ratio of growth rates (in h^-1^) for vir-recoded vs HEG-recoded variants in the indicated growth medium. Raw growth rate values for all fluorescent proteins in all media are shown in Figure S9.

### Mouse infection experiments

A schematic illustration of mice infection procedure is shown in Figure 5e. Briefly, C57BL/6J Slc11A1^r/r^ mice (21) were co-infected 1:1 by tail-vein injection of 100 μL phosphate-buffered saline containing 1,000–2,500 CFU *Salmonella* (per strain) grown to late-log phase in Lennox LB for each bacterial strain. Inoculum size was determined by plating for each infection. We used both female and male mice. Mice were scored daily for disease severity using predefined criteria covering spontaneous behavior, provoked behavior, physical appearance, clinical signs, hydration status, and grip strength. Six days after infection, animals were euthanized with CO2 and spleens were collected. Bacterial load was quantified by plating on LB agar and bacterial strains were discriminated using flow-cytometry, based on the constitutive fluorescence reporter cassette. We used *mNeonGreen* and *ypet* to discriminate vir- and HEG-recoded *mtagBFP2* expressing strains and we used *mCherry* and *mtagBFP2* to discriminate vir- and HEG-recoded *ypet* expressing strains. Competitive index (CI) values were determined as the output bacterial ratio (vir-recoded/HEG-recoded) divided by the inoculum ratio (vir-recoded/HEG-recoded): 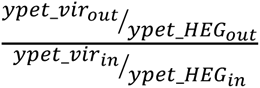. All animal experiments were approved by the Kantonales Veterinäramt Basel (license 2239) and were performed in accordance with local regulations and Swiss animal protection law. Mice were housed at 22°C (−2/+3°C), 55% ± 10% relative humidity, and a 12 h/12 h light/dark cycle. Animals were infected at 10–15 weeks of age. Sample size was estimated using a sequential statistical design. Effect size and variance were based on previous studies (15,21), and final group sizes were adjusted to achieve adequate statistical power. Experiments were not randomized.

### Computational analysis of codon usage, clustering, and sliding-window functional enrichment

All genomes CDS FASTA files were obtained from NCBI RefSeq and referenced in Table S1. For codon usage clustering analysis and more advanced computational analyses performed in this work, we developed a custom Python workflow named CodonPipe. CodonPipe is distributed through a graphical user interface and is available at Zenodo (https://doi.org/10.5281/zenodo.20486502). For each coding sequence, the pipeline computes absolute codon usage, synonymous relative codon usage, and genome-centered Z-scores of synonymous codon usage. For *STm*, the resulting codon usage tables are provided in Data S3. Because clustering based on absolute codon usage or amino acid composition is dominated by membrane-protein amino acid biases, all main analyses were performed using genome-centered synonymous codon-usage Z-scores. Practically, stop codons were excluded from analysis, synonymous codon usage was computed and Trp / Met unique codons were set to a value of 1. In the case a codon was absent from an entire cds, its value was set as undefined in exported tables and cluster averages, but for dimensionality reduction, it was replaced by the corresponding genome-wide average. Dimensional reduction was applied over the entire genome, on a genes x codons matrix. UMAP parameters were set to neighbors = 10, min dist = 0.01, cosine distance, 2 components and t-SNE parameters were set to perplexity = 10, distance = cosine, dimensions = 2, exaggeration = 10 and learning rate = 100. Genes and codons were further partitioned by k-means clustering (K = 12, ordering metric = euclidean) and then ordered within clusters by similarity to generate a one-dimensional gene order for codon-by-gene heatmaps as shown in Figure 2a.

To easily identify functional clusters, we implemented a sliding-window functional enrichment analysis along the reordered genome. Consecutive windows of 100 genes, advancing in steps of 25 genes, were submitted to DAVID (22) through the web service using mapped locus tags. For each window, the top three DAVID annotation clusters were recorded with the corresponding enrichment scores and p values. Clusters were further refined based on literature and genomic architecture (in particular pathogenic islands, plasmids and prophage regions). Resulting functional gene clusters are provided as a supplementary table for a list of species that were analyzed in this work, including *B. pseudomallei*, *B. subtilis*, *E. coli* EHEC O157:H7, *E. coli* UPEC UTI89, *K. pneumoniae*, *L. monocytogenes*, *P. aeruginosa* PAO1, *S. cerevisiae*, *S. enterica* SL1344 and *S. flexneri* 2a (Data S1). P values were plotted along the reordered genomic axis to highlight functional enrichment as shown for *Salmonella* in Figure S4. Next, user-defined gene clusters were projected onto the 2D UMAP space (*e.g.* Figure 3 for *STm*) and along the UMAP-reordered genome as flattened density plots (*e.g.* Figure 2b for *STm*).

To reduce the risk that biological interpretations depended on dimensional reduction/clustering artifacts that can arise from nonlinear methods such as UMAP and t-SNE, all dimensionality-reduction analyses were repeated using fixed random seeds and a range of UMAP and t-SNE parameters. Cluster assignments and functional enrichments were considered robust only when the same biological groups were recovered across independent seeds and parameter ranges. In addition to visualization in 2D embeddings, pairwise separation of gene sets was quantified using the Fasano–Franceschini two-dimensional Kolmogorov–Smirnov test, followed by Benjamini–Hochberg correction for multiple comparisons.

### Identification of virulence genes and definition of virulence clusters using ABRicate and Kleborate

To enhance coverage and specificity for identification of virulence genes in bacterial genomes, we used ABRicate (23) against curated virulence-factor databases. All genomes were screened against VFDB (24) and VICTORS (25), and *Escherichia coli* & *Shigella flexneri* genomes were additionally screened against the VirulenceFinder database (26) to improve detection of pathotype-specific virulence determinants. Hits were retained using minimum thresholds of 80% nucleotide identity and 60% sequence coverage. For *K. pneumoniae*, we used Kleborate to identify species-specific virulence loci (27). Hits from the different databases were merged and redundant entries were eliminated. Next, gene hits were manually verified to eliminate false positives, mainly housekeeping genes and stress response genes (H-NS, cold and heat shock proteins, ATP synthase complex, RecA, GyrA, SpoT, Lon, RpoS, ClpP,…). Manually curated tables were further grouped into functional categories, including T3SS, T6SS, LPS / O-antigen / capsule, toxins, adhesins / fimbriae / pili / curli, siderophores / iron uptake and flagella / motility / chemotaxis. Functional categories can be found in Data S1 and served as input for downstream codon decoding and tRNA usage analyses in the case of *STm*, EHEC O157:H7, UPEC UTI89, *S. flexneri* 2a and *K. pneumoniae*. Virulence genes subclusters were finally annotated using the respective genome annotation files. Annotated virulence clusters can be found at Data S4 for *STm*, EHEC O157:H7, UPEC UTI89, *S. flexneri* 2a and *K. pneumoniae*. The entire virulence discovery and clustering pipeline was generated in Python and is available at Zenodo (https://doi.org/10.5281/zenodo.20487558).

### Definition of starvation insensitive codons and genes

Starvation insensitive codons were directly extracted from (11) using as criterion that codons sensitivity as defined by the authors should be < 4 (relative units). This yielded the following list of insensitive codons: leu UUG/UUA, arg AGG/AGA/CGG, Ile AUA, Thr ACG, Ser AGU/AGC, ala GCG/GCA/GCU, Gly GGG/GGA, pro CCC/CCU. Insensitive genes as represented in flattened density plots (*e.g.* Figure 2b for *STm*) were defined as the top 10% genes in terms of enrichment in insensitive codons.

### Computational analysis of decoding strategies

Codon decoding and tRNA usage analyses were performed with the decoding analysis module of CodonPipe using a custom codon–anticodon table for *E. coli* (Table S5). tRNA abundance values were extracted from (28). To enhance analysis specificity, virulence clusters were restricted to 2D UMAP cluster localization. These refined virulence clusters were named “Vir cluster” in Data S1. To avoid assigning uncertain fractional decoding weights when a codon could be read by more than one tRNA isoacceptor, anticodons within the same amino acid family were grouped into pooled decoding units whenever individual tRNAs could not be uniquely assigned to specific codons. Accordingly, codons were mapped either to single anticodons or to multiple pooled anticodons such as UCU/CCU, UUG/CUG, or UAG/GAG/CAG. These grouped anticodons were then treated as the fundamental decoding units in subsequent analyses.

Watson–Crick versus wobble preferences were computed directly from relative codon usage z-scores (or ZCU) for amino acids associated with a unique tRNA and two synonymous codons (Tyr, Phe, Asp, Asn, His, Cys, Glu and Lys). tRNA usage was computed by summing codon counts across all codons assigned to each single or pooled tRNA(s), followed by calculation of (i) absolute tRNA usage, (ii) relative tRNA usage within a tRNA isoacceptor group (equivalent to synonymous codon usage), and (iii) genome-centered z-scores of relative tRNA usage (or ZTU). This approach avoided imposing arbitrary decoding processivity per tRNA isoacceptor, which remains unknown to our knowledge, even in *E. coli*. This simplification nevertheless preserved robust comparisons among gene clusters.

### Cross-species quantification of virulence-associated decoding signatures

To quantify cross-species decoding signatures (Figure 6e), gene-level relative codon usage values were extracted separately for ribosomal-protein genes and virulence genes in each pathogen. Enrichment scores were calculated at the gene level using relative codon usage values. For each predefined codon set, the relative usage of selected codons was summed within each synonymous amino-acid family and then averaged across amino-acid families, thereby reducing confounding by amino-acid composition. Wobble-decoded codon enrichment was calculated using U-ending codons from two-codon amino-acid families, excluding Glu and Lys because these codons were not differentially used between ribosomal and virulence gene clusters (Figure 4a). Enrichment in starvation-insensitive codons was calculated from a predefined codon set comprising Leu UUG/UUA, Arg AGG/AGA/CGG, Ile AUA, Thr ACG, Ser AGU/AGC, Ala GCG/GCA/GCU, Gly GGG/GGA, and Pro CCC/CCU extracted from (11). Enrichment in the 10% rarest tRNAs was calculated from codons decoded by the least abundant tRNA groups, using tRNA abundance values extracted from Table S5 (28). Because relative codon usage reports synonymous codon choice, rare tRNA groups decoding all synonymous codons of an amino acid, or single-codon amino acids, were excluded from this score. Usage of virulence-associated tRNA groups was calculated from codons decoded by Arg-CCG, Arg-UCU/CCU, Gly-UCC/CCC, Ile-CAU, and Leu-UAA/CAA tRNA groups, as these were significantly enriched in virulence genes in Figure 4b. Ribosomal-preferred and ribosomal-avoided codons were defined independently for each species as codons whose mean relative usage in ribosomal-protein genes was respectively above or below the uniform synonymous expectation within each amino-acid family. Gene-level scores were compared between virulence genes and ribosomal-protein genes using Welch’s t-test. Separation between virulence and ribosomal genes in codon-usage space was quantified using the regularized Mahalanobis distance between UMAP centroids, normalized by pooled within-cluster covariance, with empirical p values obtained by permutation of cluster labels.

### Statistical Analyses

Statistical significance of cluster localization along the codon-usage heatmap (*e.g.* Figure 2b) was assessed using a one-dimensional Kolmogorov–Smirnov test. For each gene cluster, the distribution of gene positions along the reordered heatmap axis was compared with the null hypothesis of a uniform distribution. Significance levels were indicated on the corresponding density plots using the conventional star annotation system.

Differences between 2D distributions of gene clusters in UMAP (or tSNE) space (*e.g.* Figure 3) were assessed using the Fasano–Franceschini two-dimensional Kolmogorov–Smirnov test. For each comparison, genes were represented by their coordinates in the 2D space obtained by UMAP or t-SNE analysis. The test was applied to compare the spatial distribution of each gene cluster against each other and against the entire genome. When multiple pairwise comparisons were performed, resulting P values were corrected for multiple testing using the Benjamini–Hochberg procedure. Corrected significance values were used to identify gene clusters showing significantly different distributions in the 2D codon-usage space. This analysis is available directly from CodonPipe.

For reporter fluorescence and infection experiments, effect sizes are reported together with exact P values and replicate numbers. One-tailed tests were chosen a priori because the reporter design tested a directional prediction, as described hereafter: (i) for mouse infection experiments (Figure 5f), we used a one-sample Student’ t-test of log10(CI) against 0, corresponding to equal fitness of vir- and HEG-recoded strains. (ii) For in vitro fitness values (Figure 5f, Figure S9), we used a two-tailed Student’ t-test to compare vir- and HEG-recoded strains. No additional log-transformation was applied, because fitness values were extracted from growth rates determined from log-transformed growth curves. (iii) For vir/HEG fluorescence ratio comparisons (Figure 5c), vir/HEG fluorescence ratios measured in the absence or presence of casamino acids were compared using a one-tailed paired Student’s t-test, with the alternative hypothesis that vir/HEG ratios are higher under starvation/no-casamino acid conditions than in the corresponding casamino acid-supplemented condition. (iv) Codon, tRNA and amino acid usages (Figure 4, Figure S2c, Figure S8, Figure S12) were tested using two-tailed Student’s t-tests.

## Results

### CodonPipe reveals unexpected complex and structured codon usage landscapes across bacteria

To visualize codon usage patterns in bacterial genomes, we developed **CodonPipe**, an integrated platform for codon usage profiling based on dimensional reduction and clustering to reorder genes according to codon similarity and codons based on their cooccurrence across the genome. The framework thus generates as output genome-level heatmaps, codon usage comparisons across sets of genes, functional enrichment analyses and other features explained further in this manuscript. A schematic view of CodonPipe is shown in Figure 1.

**Figure 1.**
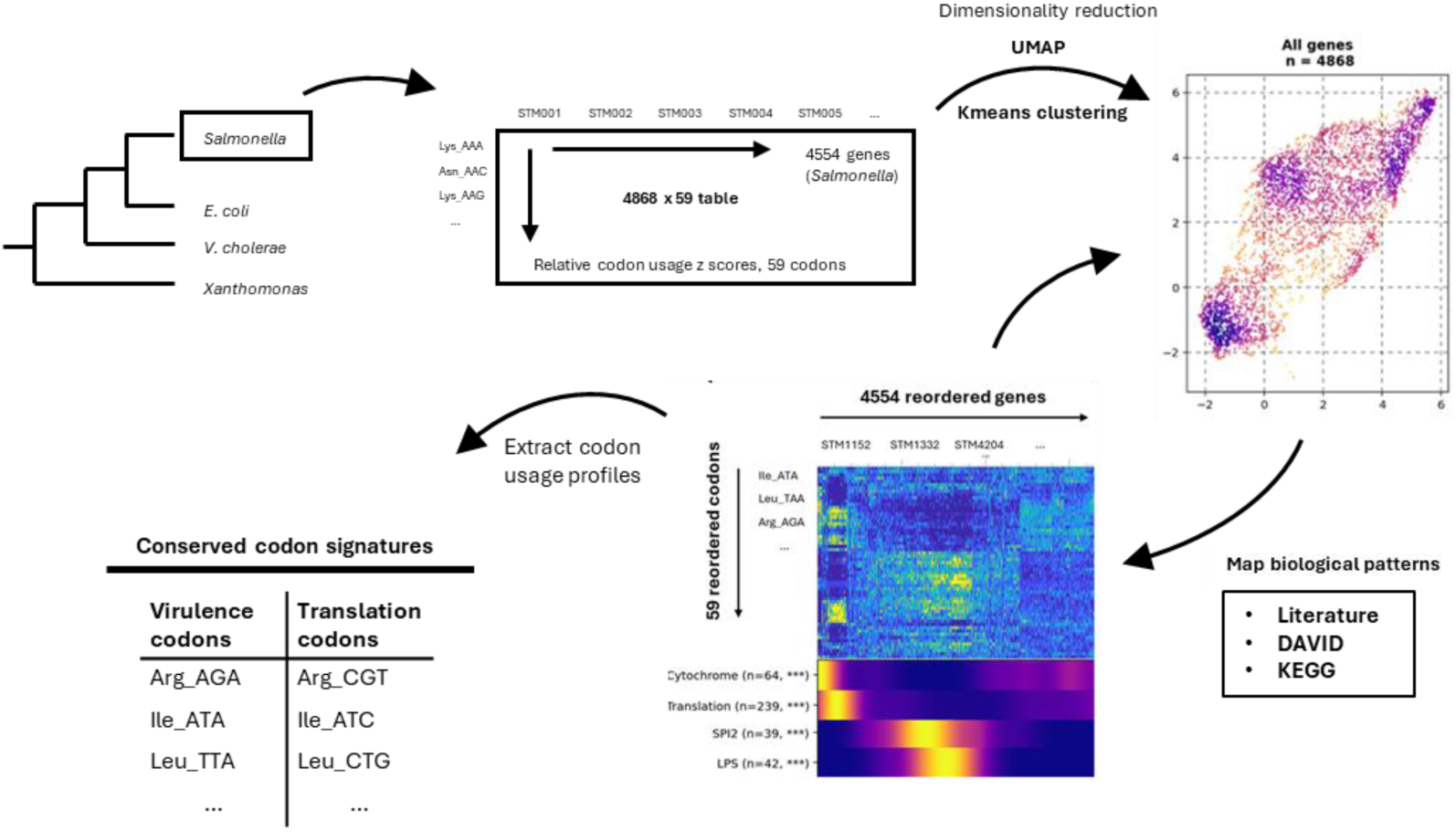
CodonPipe workflow. Coding sequences are converted into synonymous codon-usage profiles, normalized as genome-centered codon-usage Z-scores, represented in a 2D UMAP/t-SNE/PCA space, clustered, and reordered to generate genome-wide codon-usage landscapes. Functional gene sets are then mapped as density maps onto the reordered genomic axis and into the 2D UMAP/t-SNE/PCA space.

We first employed the widely used principal component analysis (PCA), a linear clustering method. PCA separated some major gene classes in the model organism *Salmonella enterica* serovar Typhimurium SL1344 (further *STm*), including ribosomal and virulence genes (1), but provided limited overall resolution (Figure S1a). We then tested non-linear dimensionality reductions with Uniform Manifold Approximation and Projection (UMAP) (29), yielding a rich and well-resolved landscape of unique codon usage patterns (Figure 2a). Such rich patterns were robust against variation of critical parameters such as number of neighbors and minimal distance (Figure S1b,c). A similarly rich and robust pattern emerged using another non-linear dimensionality reduction method, t-distributed stochastic neighbor embedding (t-SNE) (Figure S1d,e). In both cases, AT bias and gene origin (ancestral vs horizontal) contributed (Figure 2b, three first rows), but did not fully explain the patterns, while chromosome-strand biases in codon usage had little impact (Figure 2b, fourth row). We found that clustering based on absolute codon frequencies (Figure S2a) or amino acid composition (Figure S2b) revealed large and distinct clusters of inner and outer membrane proteins. For inner membrane proteins, this was consistent with significant enrichment in hydrophobic and aromatic residues Leu, Ile, Met, Phe and Trp (Figure S2c). For outer membrane proteins, Asn, Asp, Ser, Thr and Tyr residues were significantly more abundant, likely reflecting abundant β-barrel structures in outer membrane proteins (Figure S2c). These biases were prevented by clustering according to genome-centered synonymous codon-usage z-scores (further ZCU) (Figure S2d).

**Figure 2.**
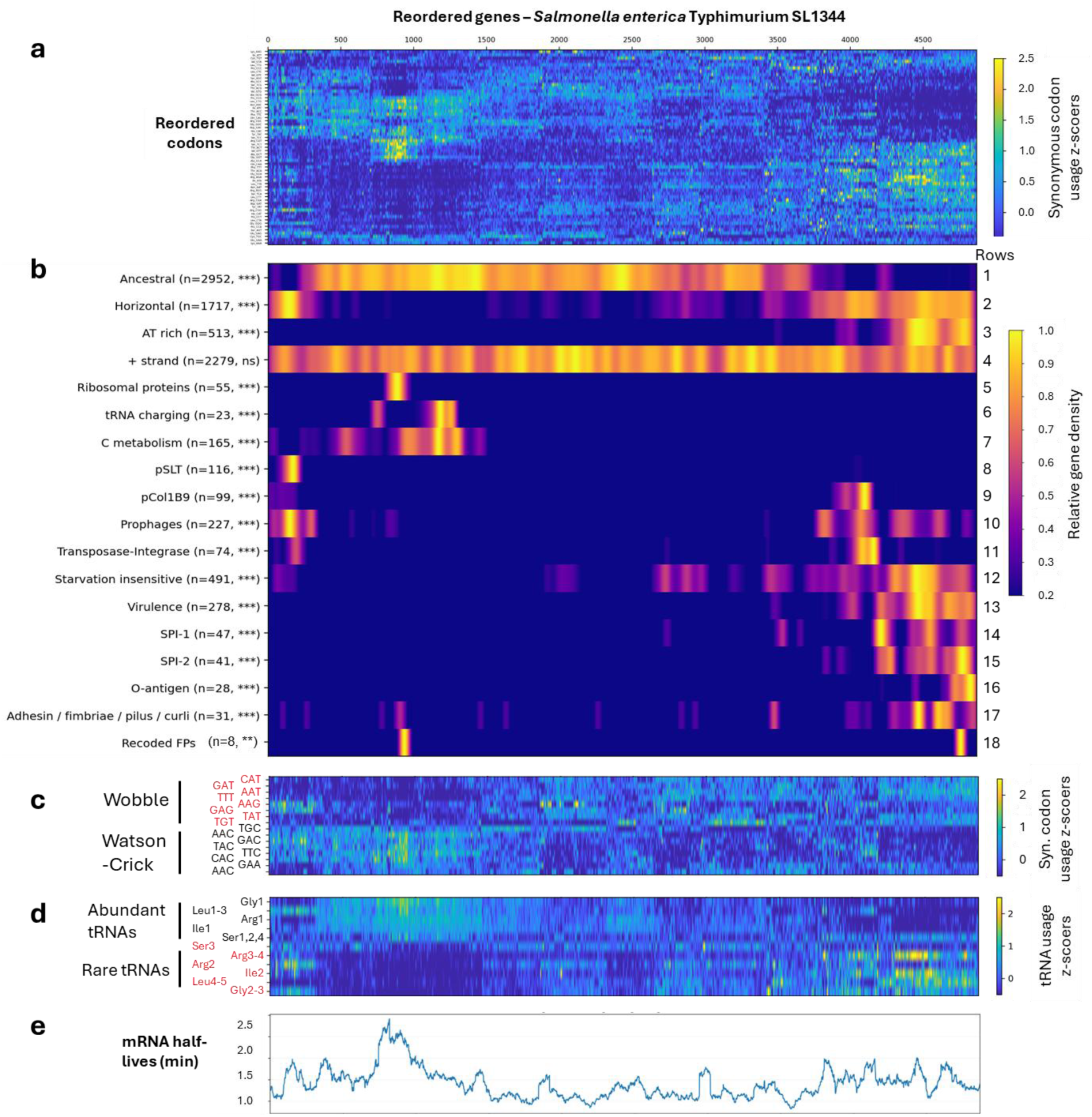
UMAP clustering reveals diverse, functionally distinct codon usage signatures in *STm*. (a) Heatmap of synonymous codon-usage Z-scores (ZCU) after genes and codons were reordered by codon-usage similarity using non-linear dimensionality reduction (UMAP) followed by k-means clustering. (b) Density enrichment of major gene clusters along the reordered genomic axis. Gene counts are indicated. Non-random localization was assessed by one-dimensional Kolmogorov-Smirnov tests. (c) Wobble versus Watson-Crick decoding preference across the reordered gene axis for amino acids decoded by a single tRNA species. (d) tRNA-usage Z-scores (ZTU) across the reordered gene axis for amino acids decoded by multiple tRNA isoacceptors. (e) mRNA half-life estimates from (30) plotted across the reordered genomic axis and smoothed with a 75-gene running average.

Across 28 bacterial species and the yeast *S. cerevisiae*, spanning GC content from 32% (*L. monocytogene*s) to 68% (*C. crescentus*) (Table S1), codon usage landscapes were similarly rich, although the specific signatures differed (Figure S3). Genome size could not account for this diversity, as even the small genomes of *S. pneumoniae* (2116 genes) and *S. pyogenes* (1649 genes) harbored complex codon usage landscapes. Only *Helicobacter pylori* and *Francisella tularensis* showed less clearly resolved patterns. This is consistent with small AT-rich genomes, extensive recombination or horizontal gene transfer in *H. pylori* (31) or reductive adaptation to intracellular niches with enrichment in pseudogenes and IS elements in *F. tularensis* (*32*). Strikingly, *H. pylori* and *F. tularensis* encode unusually small tRNA sets (36–38 genes vs 86 *in E. coli*), consistent with reduced codon diversification. The absence of codon clustering in these pathogens therefore supports the validity of our approach.

Thus, CodonPipe revealed reproducible, functionally structured codon-usage landscapes in most genomes analyzed. Because nonlinear embeddings can exaggerate local structure, we treated UMAP and t-SNE primarily as visualization and ordering tools, and interpreted clusters only when they were supported by enrichment analysis, robustness across parameters, and separation in the original synonymous codon-usage feature space. This conservative strategy allowed us to move from descriptive codon landscapes to testable biological hypotheses.

### Diverse cellular functions adopt distinct codon signatures, extending beyond previous knowledge

Non-linear dimensionality reduction methods such as UMAP and t-SNE can be superior to linear PCA to reveal structure in complex or non-linear data, curved manifolds, and data with high local densities. However, UMAP and t-SNE do not preserve distances or densities of the original data and can artificially create more clusters than genuinely exist (33–35). Thus, it is crucial to determine if structures revealed by non-linear methods may have biological significance. To facilitate this, we implemented an unbiased functional enrichment analysis scanning the codon-usage landscape with a sliding window of 100 genes advancing by steps of 25 genes across the entire codon-usage landscape (see methods). This yielded a series of functional enrichment peaks along the reordered genome axis as shown for *STm* as an example (Figure S4). Clusters were further refined based on literature and genomic architecture (pathogenic islands, prophages, plasmids). In *STm*, multiple functional clusters formed distinct distributions within the 2D UMAP space and along the UMAP-reordered genome (Figure 2b, Figure 3). This included a tight, well-known cluster of ribosomal proteins and another cluster of virulence-associated genes dominated by *Salmonella* type III secretion systems (T3SS) and O-antigen biosynthesis genes. Genes associated with carbon metabolism and tRNA acylation formed two additional clusters close to but not overlapping with ribosomal genes. Horizontally acquired genes were defined by 3 major clusters corresponding to (i) prophage and plasmid genes, (ii) transposase and integrase genes and (iii) pathogenicity islands, largely overlapping with the virulence cluster.

Statistical analysis confirmed that clusters were distinct from one another (2D Kolmogorov-Smirnov, BH-adjusted for multiple comparisons, Table S2) but also revealed the existence of subclusters. In particular, virulence genes formed four separate sub-clusters of genes associated with (i) the cell invasion-linked type 3 secretion system 1 (T3SS-1), (ii) the intracellular growth-linked T3SS-2, (iii) the biosynthesis of the O-antigen of LPS and (iv) adhesins, fimbriae and curli (Figure 2b rows 13-17, Figure 3). Conversely, *Salmonella* type VI secretion system did not show distinct codon-usage clustering and did not colocalize with the virulence cluster. In addition, there were multiple other codon usage signatures, but they showed no enrichment for particular gene ontology terms, suggesting that they might be more complex modules involving multiple gene functions.

**Figure 3.**
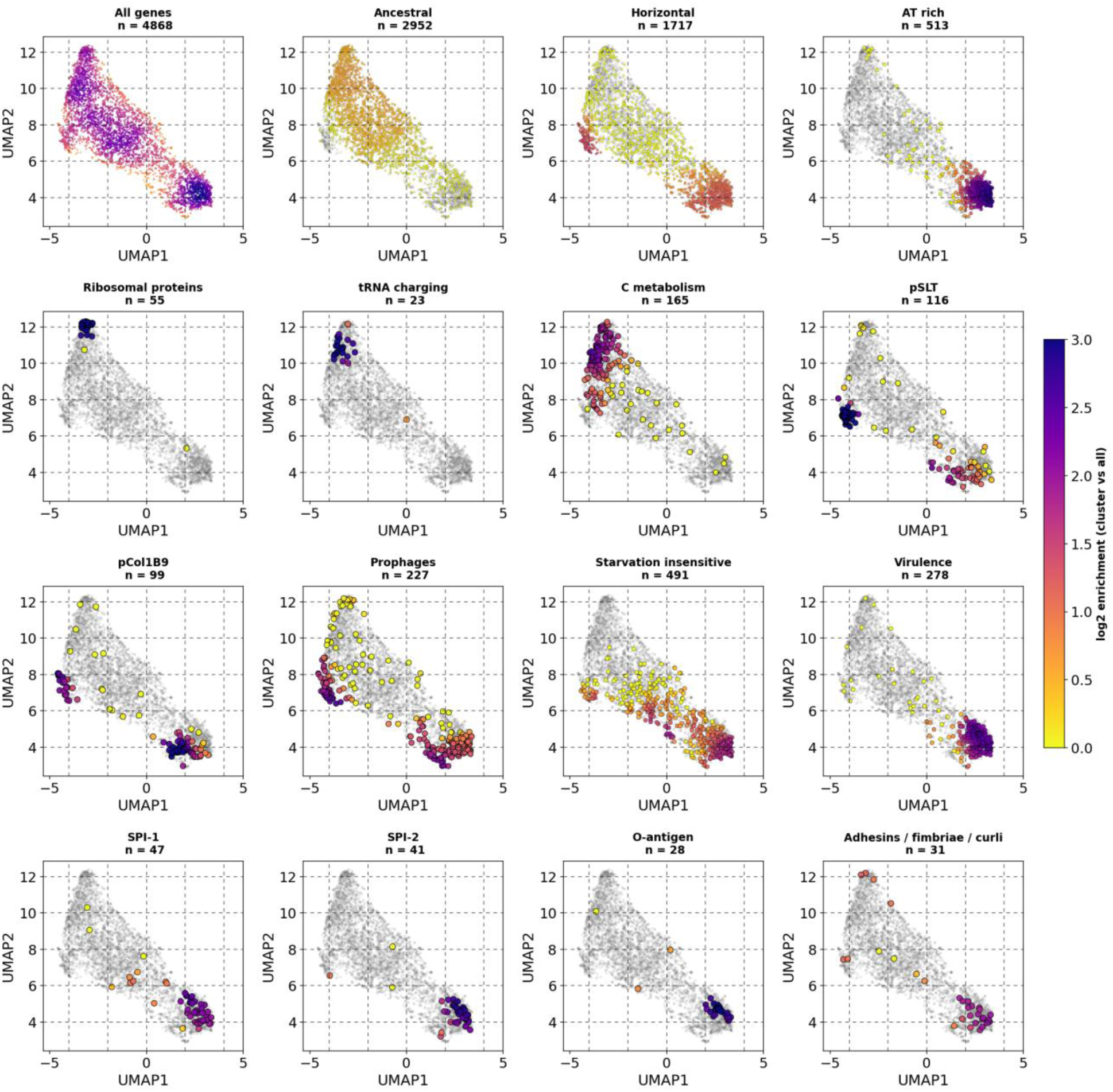
Functional gene clusters occupy distinct distributions in the *STm* codon-usage space. 2D UMAP plots show all coding sequences in grey and density enrichment for representative functional gene clusters. Color intensity indicates local enrichment relative to the whole-genome background. Gene counts are indicated above each panel.

Intriguingly, *STm* plasmids pSLT and pCol1B9 as well as prophages, each separated into two distinct subclusters, one overlapping with the virulence cluster and the other positioned closer to the ribosomal genes cluster (Figure 3, Figure S5, data S5). A similar organization was also observed for the *E. coli* F plasmid (Figure S6, data S5). The subcluster closer to ribosomal genes was enriched in plasmid maintenance, establishment and transfer genes, whereas the virulence-associated subcluster showed hallmarks of cargo/accessory genes (36), including the *spv* and *pef* loci, which contribute to systemic virulence through the T3SS-2 translocated effectors and to *Salmonella* adhesion, respectively. However, we could not attribute this clustering to distinct phylogenetic origins co-existing within plasmids, as illustrated by conjugation machinery genes that were broadly distributed across both subclusters.

We next extended this analysis to *Listeria monocytogenes*, *Pseudomonas aeruginosa*, *Burkholderia pseudomallei*, *Bacillus subtilis* and *Saccharomyces cerevisiae*. In all genomes, we identified multiple functional clusters with distinct codon-usage signatures, including ribosomal proteins, aminoacyl-tRNA synthetases, carbon metabolism genes, transposases and integrases, prophages and plasmids as well as virulence genes in pathogenic species (Figure S7 a-e). Species-specific clusters were also detected, including internalins in *L. monocytogenes*, porins in *B. pseudomallei*, diverse phage-associated signatures in *B. subtilis* and Ty LTR retrotransposon, ATP-dependent helicases and SRP1/TIP1 genes in *S. cerevisiae*. Cluster compositions for all species are listed in Data S1.

These observations argue against a purely phylogenetic explanation for the codon-usage landscape. Horizontally acquired regions did not collapse into a single signature; instead, prophages, plasmids, transposases and virulence-associated loci occupied partially distinct regions of codon space. Conversely, functionally related virulence genes converged toward similar codon signatures despite differing genomic locations and evolutionary histories. We therefore asked whether this organization reflected a shared decoding strategy rather than only a shared origin.

### Virulence codons avoid direct competition with highly expressed genes by favoring rare tRNAs and wobble decoding

To identify some of the drivers of this remarkable codon usage complexity, we compared codon choice across major *STm* gene clusters. Ribosomal genes, which strongly deviated from the genomic average (Figure 2b row 5), showed the expected hallmarks of translation optimization. Notably, this signature was not associated with higher GC content: ribosomal genes had a slightly lower GC content than the overall *STm* genome (51.7% versus 53.4%). For amino acids with synonymous codons decoded by a single anticodon (Asn, Asp, Cys, Glu, His, Lys, Phe, and Tyr), ribosomal genes preferentially used codons decoded through canonical Watson–Crick base pairing rather than wobble interactions (Figure 2c, Figure 4a). This is consistent with increased codon-anticodon stability and minimization of translational errors (missense/nonsense), thereby limiting accumulation of misfolded proteins that can cause pleiotropic toxicity in bacteria (37,38). Lys was an exception, showing no stronger preference for AAA over AAG (Figure 4a), possibly because the genome already exhibits an approximately 3:1 bias toward AAA. It was intriguing that for Tyr, Phe, His and Asp, Watson-Crick decoding was not the first choice at the genome level. This suggests additional selective mechanisms maintain wobble decoding for specific amino acids, despite its potential cost in translational accuracy.

We next sought to investigate tRNA choices. This can be challenging because several codons can be decoded by multiple tRNAs either through Watson-Crick or wobble interactions and the fraction decoded by each tRNA remains unknown. To circumvent this limitation, we grouped tRNA isoacceptors when necessary, so that codons could be assigned unambiguously to defined decoding units. In *STm*, ribosomal genes systematically favored the most abundant tRNAs (Figure 2d, Figure 4b), consistent with previous observations (39,40).

**Figure 4.**
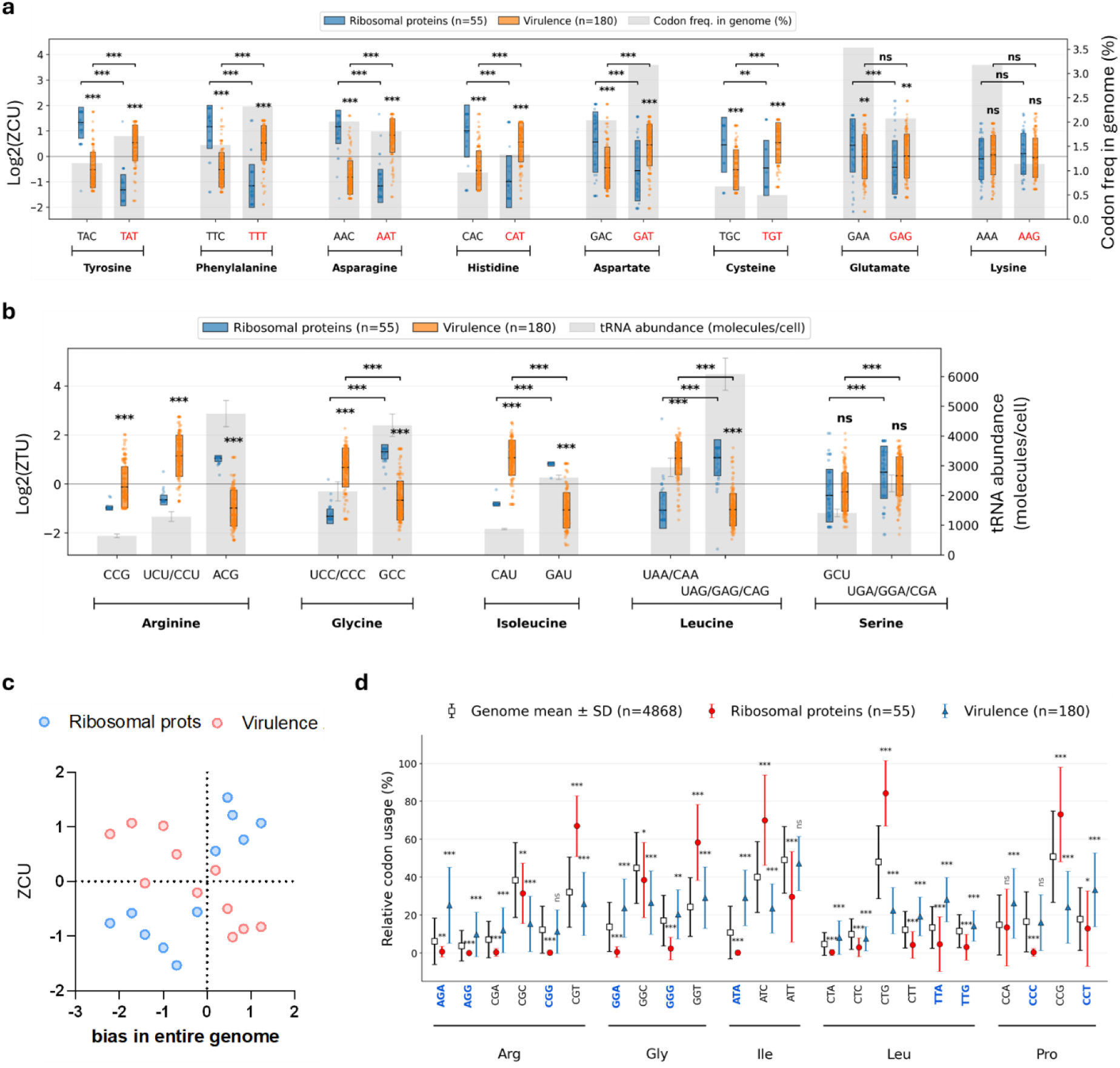
*Salmonella* virulence genes use decoding strategies that avoid competition with ribosomal genes and enable robust expression during starvation. (a) Synonymous codon-usage Z-scores for codons decoded through Watson-Crick or wobble pairing in amino acid families decoded by a single tRNA. Grey bars indicate genome-wide codon frequencies in *STm*. (b) tRNA-usage Z-scores for amino acids decoded by multiple tRNA isoacceptors, shown either individually or pooled into isoacceptor groups when codons could not be unequivocally assigned. Grey bars indicate tRNA abundance from (28). (c) Mean codon-usage shifts in ribosomal versus virulence genes versus average genomic codon usage in *STm*. (d) Relative usage of starvation insensitive codons (in blue) as defined by (11,12). Data are means ± SD. In panels a, b and d, statistical significance was assessed using two-tailed Student’s *t*-tests: * *P* < 0.05, ** *P* < 0.01, *** *P* < 0.001.

Virulence genes showed the opposite tendency (Figure 2b row 13). For Asn, Asp, Cys, His, Phe, and Tyr, they preferentially used U-ending codons decoded by wobble interactions rather than C-ending codons decoded by canonical Watson–Crick pairing (Figure 4a). Although this was partly consistent with the lower GC content of virulence genes compared with the overall *STm* genome (45.8% versus 53.4%), the pattern could not be explained by A/T enrichment alone. In particular, for Glu and Lys, where A-ending codons correspond to Watson–Crick decoding, virulence genes showed no additional enrichment in A-ending codons. Instead, virulence codon choices were inversely related to those of ribosomal genes (Figure 4c), both at the codon level (P = 0.048) and even more strongly at the level of tRNA isoacceptor choice (P = 0.0047). This inverse relationship was associated with increased usage of rare tRNA isoacceptors for Arg, Gly, Leu and Ile (Figure 2d, Figure 4b). Thus, virulence genes did not simply favor A/T-rich codons but adopted a decoding strategy opposite to that of highly expressed ribosomal genes.

Because codon usage can also influence mRNA stability (6–8), we next asked whether the unusual codon usage of virulence genes was associated with destabilized transcripts. To address this, we used a recent transcriptome-wide analysis of mRNA stability in *Salmonella*, inferred through Bayesian modeling (30), and projected these values onto the UMAP-reordered genome (Figure 2e). As expected, ribosomal genes coincided with a major peak of transcript stability. However, virulence genes were also associated with stable transcripts compared with the rest of the genome, indicating that their atypical codon usage does not cause broad transcript destabilization.

Together, these findings challenge the common view that low GC content and atypical codon usage represent an intermediate evolutionary state of recently horizontally acquired genes undergoing adaptation to the host genome (2,4). Instead, our data suggest that virulence genes may actively exploit this decoding strategy to avoid direct competition with highly expressed genes while preserving transcript stability. This strategy may be particularly important during host infection, when bacteria must sustain bursts of virulence gene expression while facing host-imposed stresses, including nutrient deprivation. Because tRNA isoacceptor charging levels vary strongly during amino acid starvation, thereby explaining the enrichment of rare codons in genes encoding amino acid biosynthetic enzymes (11,12), we asked whether a similar mechanism could support selective translation of virulence genes under host-imposed nutrient starvation.

### Virulence codon bias promotes robust translation in *Salmonella* during stress and starvation

tRNAs heavily used for translation of ribosomal genes are particularly sensitive to sudden amino acid starvation, because their aminoacylated forms are rapidly consumed by abundant transcripts but cannot be efficiently recharged when amino acids become limiting (11,12). Based on prior work that quantified sensitivity of synonymous codons to amino acid starvation, we defined starvation-insensitive codons using published sensitivity scores that were based on mathematical modeling (11) and were experimentally validated using microarrays for measuring charged fractions of tRNAs during leucine, threonine and arginine starvation in *E. coli* (12). In contrast to ribosomal genes, we found that virulence genes preferentially used insensitive codons, including Arg AGA/AGG/CGG, Gly GGG/GGA, Ile ATA, Pro CCC/CCT and Leu TTA/TTG (Figure 4d). We next identified the top 10% of genes most enriched in these codons in *STm* genome and found that they closely recapitulated the overall codon usage profiles, tRNA isoacceptor choice and decoding strategy of virulence genes, but not those of ribosomal or carbon metabolism genes (Figure 2b row 12; Figure S8a,b). More surprisingly, they also differed from prophage and plasmid genes. Although plasmid and prophage signatures partially resembled those of virulence and starvation-insensitive genes, they avoided Leu TTA/TTG codons decoded by rare tRNA^leu^ ^UAA/CAA^, similarly to ribosomal genes (Figure S8c). This suggested that virulence codon usage may uniquely help preserve translation under nutrient stress, including during invasion through body surfaces or entry into host cells, which represent vulnerable steps in disease progression.

To test this model experimentally, we generated fluorescent protein variants recoded to match either ribosomal/highly expressed genes (HEG) or virulence genes (vir) codon usage (Figure 5a). These variants were expressed in *STm* either constitutively from ProD (*gfpmut2*) or from an arabinose-inducible pBAD promoter (*mtagBFP2* and *ypet*). To minimize differences in mRNA stability, the first 25 codons were kept identical between HEG- and vir-recoded versions of each gene (6–8). In addition, all fluorescent proteins carried a C-terminal LVA degradation tag (19), limiting potential differences caused by protein accumulation or sequestration of amino acids. As expected, HEG- and vir-recoded genes mapped onto the UMAP-reordered *STm* genome with ribosomal and virulence gene clusters, respectively (Figure 2b, row 18). Their codon composition also shifted as intended, from 31–33% to 23–26% starvation-sensitive codons and from 4–6% to 14–19% starvation-insensitive codons in HEG- and vir-recoded fluorescent proteins, respectively (Figure 5b). As a control, we verified that expression of the recoded fluorescent proteins did not detectably impair growth under the tested in vitro conditions (Figure S9).

**Figure 5.**
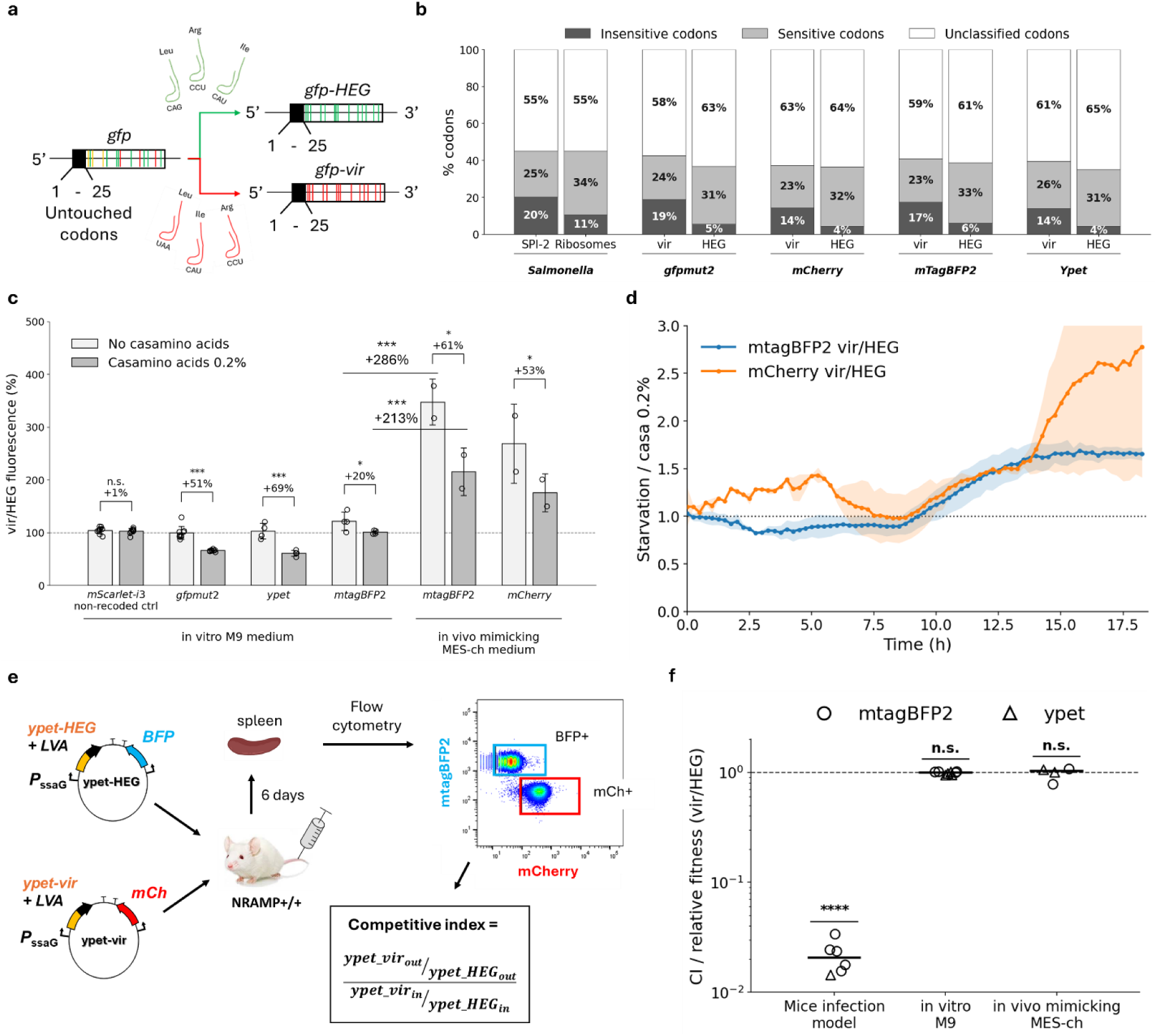
Virulence codon bias promotes robust translation in *Salmonella* during stress and starvation. (**a**) Design of paired fluorescent-protein reporters. For each pair, the first 25 codons were kept identical, whereas the remaining coding sequence was recoded to match either ribosomal/highly expressed genes (HEG) or virulence genes (vir). (**b**) Distribution of sensitive and insensitive codons as defined in (11), in SPI-2 genes, ribosomal genes and recoded fluorescent protein coding sequences. (**c**) Vir/HEG fluorescence ratios for recoded fluorescent protein pairs grown in M9 medium or in MES-ch medium, with or without 0.2% casamino acids. Exponentially growing bacteria were diluted into 1 mL of the indicated medium in 24-well microplates, and OD_600_ and fluorescence were monitored every 15 min for 18 h. Bars show the mean vir/HEG fluorescence ratio over the analysis window defined in the methods section. Values above 100% indicate a positive effect of virulence codon usage on expression levels, as compared to ribosomal codon usage. Error bars are SD, and dots represent biological replicates (n = 2 in MES and n ≥ 3 in M9). Statistical tests were paired one-tailed Student’s t-tests, comparing vir/HEG fluorescent ratios in the same medium with or without 0.2% casamino acids. (**d**) Impact of casamino acid starvation on Vir/HEG fluorescence ratios in MES-ch medium over time. Solid lines represent mean values and shaded areas represent SEM (n = 2). (**e**) Mouse co-infection scheme using *Salmonella* Typhimurium (*STm*) Sme51 strains carrying vir- or HEG-recoded fluorescent-protein reporters. Mice were co-infected with equal numbers of both strains. After 6 days, spleens were collected, bacteria were recovered, and the two competing strains were distinguished by flow cytometry using constitutive *mtagBFP2* vs *mCherry* fluorescence for recoded *ypet* or constitutive *ypet* vs *mNeonGreen* fluorescence for recoded *mtagBFP2* (see methods). Competitive index (CI) values were calculated as indicated. (**f**) Vir/HEG CI values in mice was computed as the vir/HEG ratio of live bacteria after 6 days of infection normalized by the same ratio used for infection. vir/HEG relative fitness in vitro was calculated based on growth-rate ratios in the indicated media. Circles and triangles represent *mtagBFP2* and *ypet*, respectively. Dots represent individual mice or independent biological replicates, as appropriate (n = 6). CI values and relative fitness values were tested against expected value of 1 using one-tailed paired Student t-test. CI values were compared using one-sample Student’ t-test of log10(CI) against 0. Relative fitness differences were tested using two-tailed Student’ t-test to compare vir- and HEG-recoded strains.

We first compared HEG- and vir-recoded reporters during steady-state induction in nutrient-rich conditions, using M9 supplemented with 0.2% casamino acids. Under these conditions, HEG-recoded variants produced higher fluorescence than their vir-recoded counterparts for *ypet* and *gfpmut2*, as indicated by vir/HEG fluorescence ratios below 100%, whereas *mtagBFP2* showed the opposite trend (Figure 5c). Thus, under nutrient-rich conditions, virulence codon usage did not provide a consistent translational advantage, in line with previous reports showing that codon usage has limited impact during continuous expression in rich media (1,6–8,37,38). We then imposed sudden amino-acid starvation by washing exponentially growing cells and transferring them to M9 medium lacking casamino acids. Under these conditions, *STm* maintained growth, albeit at reduced growth rates (Figure S9). However, vir-recoded reporters outperformed their HEG-recoded counterparts, resulting in vir/HEG fluorescence ratios above 100% (Figure 5c). By contrast, a non-recoded *mScarlet-i3* control showed no comparable starvation-dependent increase. Thus, virulence codon bias selectively improved translation during sudden amino-acid starvation.

We next asked whether this effect would become more pronounced under conditions that more closely mimic the intracellular environment encountered during host infection. To this end, we placed *mtagBFP2* and *ypet* recoded variants under the control of the SPI-2-inducible promoter *PssaG* and grew *STm* in MES-ch, an acidic low-nutrient medium designed to mimic the host intracellular environment, with low phosphate (113 µM), magnesium (50 µM), glucose (0.01%) and glycerol (0.02%) at pH 5.5 (Table S3) (15). As expected, *STm* displayed reduced growth rates in MES-ch, especially in the absence of casamino acids (0.14 h^-1^ versus 0.59 h^-1^ in M9 without casamino acids) (Figure S9). Under these conditions, the vir- recoded ypet reporter produced fluorescence signals too low for robust quantitative interpretation, potentially reflecting altered mRNA abundance, maturation, folding or stability under acidic low-nutrient conditions. We therefore report these data separately (Data S6) and excluded them from the main quantitative analysis. By contrast, vir-recoded *mtagBFP2* showed a strong and significant advantage over its HEG-recoded counterpart that increased over time (Figure 5d). The vir/HEG fluorescence ratio increased substantially as compared to M9 in the presence of casamino acids (+213%, P = 0.00069) and even more so in the absence of casamino acids (+286%, P = 0.00048) (Figure 5c). Moreover, amino acid starvation increased the vir/HEG ratio by 61% (P = 0.039). To validate these results, we generated vir- and HEG-*mCherry* variants under the control of *PssaG*, which reproduced the results obtained with *mtagBFP2* (Figure 5c,d).

Together, these reporter experiments support a model in which virulence-gene codon usage improves translation efficiency during sudden nutrient limitation and under host-mimicking stress conditions. Importantly, this effect was specific: virulence-like codon usage did not provide a uniform advantage in nutrient-rich medium but became beneficial when translational resources were expected to be limiting. However, recoding of fluorescent reporters outside the shared N-terminal region could still influence mRNA stability and abundance, as observed for *ypet*. To cope with this limitation, we reasoned that overexpression of vir-recoded proteins should impose a selective burden in a mice infection model. This effect should be specific to vir-recoded proteins and be absent in vitro.

To test this, we co-infected mice with *STm* strains carrying either vir- or HEG-recoded fluorescent protein constructs and measured the competitive index of vir-recoded relative to HEG-recoded strains. At 6 days post-infection, spleens were harvested, and competing strains were discriminated by flow cytometry using constitutive fluorescent markers. For competitions between strains expressing *mtagBFP2* recoded variants, vir- and HEG-recoded strains were distinguished using constitutive *mNeonGreen* and *ypet* markers. For competitions between strains expressing *ypet* recoded variants, competing strains were distinguished using constitutive *mCherry* and *mtagBFP2* markers (Figure 5e). Expression of vir-recoded fluorescent proteins selectively disrupted virulence of *STm*, resulting in a 47-fold drop in competitive index (p = 4×10^-7^), while leaving growth unaffected in vitro in M9 medium and in host-mimicking MES-ch medium (Figure 5f). These results showed that expression of virulence in *STm* relies on distinct subsets of isoacceptor tRNAs that could be depleted by overexpressing fluorescent proteins recoded to match virulence genes codon bias. This also indicated that targeting virulence tRNA pools could constitute a promising anti-virulence strategy.

Altogether, our experimental data provide strong in vitro and in vivo supportive evidence for a model in which codon bias promotes robust virulence gene expression under stress by reducing competition with highly expressed genes and by favoring usage of tRNA isoacceptor pools that remain selectively charged during sudden starvation. However, this strategy makes virulence expression dependent on a restricted subset of tRNAs that could be selectively exploited to direct anti-virulence strategies against *Salmonella* in the future. We next asked whether this principle was specific to *STm* or reflected a broader feature of bacterial pathogens.

### Conserved virulence decoding strategies suggest that stress-adapted codon usage may represent a broader principle of bacterial pathogenesis

We therefore asked whether the decoding strategy identified in STm virulence genes was conserved across other Enterobacteriaceae pathogens with distinct infection strategies. We analyzed enterohemorrhagic *E. coli* O157 Sakai, *Shigella flexneri* 2a, uropathogenic *E. coli* UTI89 and *Klebsiella pneumoniae*. Together with *Salmonella*, these species include major human pathogens prominently represented among WHO priority antibiotic-resistant bacteria and strongly affected by the global rise of antimicrobial resistance (41,42).

To systematically identify virulence-associated clusters, we screened genomes across curated virulence databases using ABRicate (23), including VFDB (24), VICTORS (25) and VirulenceFinder (26), and complemented this analysis with Kleborate for *K. pneumoniae* (27). Genes were assigned to functional categories, including T3SS, T6SS, LPS/O-antigen/capsule biosynthesis, toxins, adhesins/fimbriae/pili/curli, siderophores/iron uptake and flagella/motility. As with *STm*, functional clusters were mapped onto the 2D codon usage UMAP space and visualized as flattened density plots along the UMAP-reordered genome (Figure 6a-d, Figure S10).

**Figure 6.**
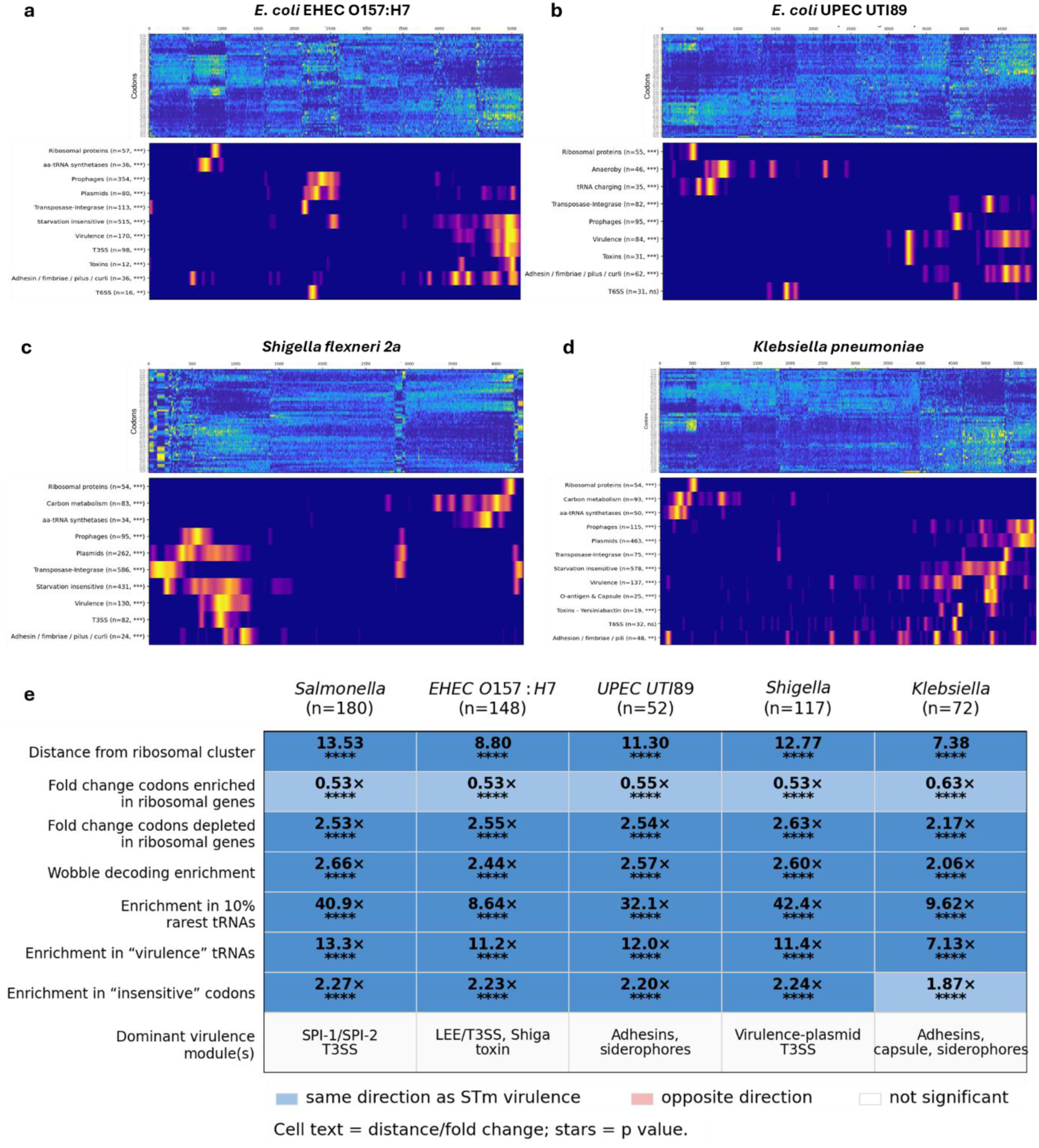
Virulence decoding signatures are shared across Enterobacteriaceae pathogens. (a-d) For each species, heatmaps show synonymous codon-usage z-scores after UMAP-based gene reordering, and adjacent density plots show enrichment of functional gene categories along the same reordered axis. Gene counts and one-dimensional KS-test significance are indicated. (e) Comparative analysis of virulence-associated decoding strategies across Enterobacteriaceae pathogens, see detailed analysis in Figure S12. Columns correspond to pathogens and rows to codon-usage or decoding metrics comparing virulence genes with ribosomal-protein genes. “Distance from ribosomal cluster” indicates the regularized Mahalanobis distance between virulence and ribosomal gene centroids in the 2D UMAP codon-usage space with significance assessed by permutation testing. “Fold change codons enriched in ribosomal genes” represents the virulence/ribosomal fold change in codons that are enriched in ribosomal-protein genes relative to uniform synonymous codon usage. “Fold change codons depleted in ribosomal genes” mirrors the previous metric for codons that are under-used in ribosomal-protein genes. “Wobble-decoded codon enrichment” represents the virulence/ribosomal fold change in usage of U-ending wobble codons, excluding Glu and Lys. “Enrichment in 10% rarest tRNAs” represents fold-change in usage of codons decoded by the 10% least abundant tRNAs as defined by (28). “Enrichment in virulence tRNAs” indicates the virulence/ribosomal fold change in codons decoded by virulence-associated tRNA groups as defined earlier (Figure 4b). “Enrichment in insensitive codons” indicates the virulence/ribosomal fold change in codons associated with robust translation during amino-acid limitation as defined by (11). For all gene-level codon and decoding metrics, p values were calculated using Welch’s t-test comparing virulence and ribosomal-protein genes. Significance is indicated as ****, p < 0.0001.

Selected pathogens covered markedly different virulence strategies (Figure 6e, Figure S11, Data S4). *Salmonella*, *Shigella* and EHEC O157 rely on distinct T3SS to drive epithelial invasion, intracellular survival or attaching-and-effacing lesions. By contrast, UPEC UTI89 and *K. pneumoniae* do not encode a T3SS and rely instead on adhesion, persistence, iron acquisition, toxins and, for *K. pneumoniae*, capsule- and LPS-mediated immune evasion. Despite these differences, virulence genes displayed strikingly conserved codon-usage organization across pathogens. In all species, virulence genes occupied regions of codon-usage space distinct from ribosomal genes and overlapped with genes enriched for starvation-insensitive codons (Figure 6e, Figure S10). This was further supported by the enrichment of virulence genes in wobble codons and “starvation insensitive” codons decoded by rare tRNAs in all pathogens (Figure 6e, Figure S12). Thus, although virulence clusters differed between species, their decoding properties converged to a common signature characterized by reduced competition with highly expressed genes and predicted enhanced translation during nutrient starvation. Note that this signature was less sharply defined in *K. pneumoniae* (Figure 6e, Figure S12j-l), likely reflecting the more heterogeneous composition of its virulence cluster and the absence of a dominant T3SS.

Altogether, these analyses indicate that codon usage signature of virulence genes is conserved across diverse Enterobacteriaceae pathogens. This supports a broader model in which virulence genes adopt decoding strategies that limit competition with highly expressed housekeeping genes while preserving robust translation under starvation, particularly during host infection. To our knowledge, this provides a first example of how host-associated environmental pressures can shape codon usage at the level of functional gene classes, even when these genes are dispersed across the genome and have diverse evolutionary origins, including core genes, pathogenicity islands, plasmids and prophages.

## Discussion

This work revisits a long-standing interpretation of bacterial codon usage. Rather than merely reflecting expression level, nucleotide composition or evolutionary origin, codon usage formed complex and structured landscapes in which genes sharing biological functions converged toward distinct decoding signatures. This organization extended far beyond the classical separation between highly expressed genes, horizontally acquired genes and the rest of the genome, suggesting that synonymous codon choice might play functional roles in the regulation of gene expression.

Virulence genes provided the clearest example of this principle. In *Salmonella*, they retained a coherent AT-rich codon signature despite diverse evolutionary origins and genomic locations. This signature could not be explained by A/T enrichment alone. Instead, we found that virulence genes exploit a decoding strategy opposite to that of highly expressed housekeeping genes, favoring wobble interactions, rare tRNAs and codons previously shown to support robust translation during amino acid starvation (11,12). These findings suggest that virulence codon usage might reduce competition with housekeeping translation and promote robust expression

Our reporter experiments support this model. Fluorescent proteins recoded to mimic virulence-gene codon usage showed a selective advantage during sudden nutrient limitation and in acidic, low-phosphate, low-magnesium host-mimicking medium. During mice co-infections, overexpression of virulence-recoded reporters imposed a selective in vivo burden relative to HEG-recoded reporters. The conservation of this signature across diverse Enterobacteriaceae pathogens further suggested that this is not specific to *Salmonella*, but may represent a broader principle of bacterial pathogenesis. Even when virulence genes differ in function, genomic location and evolutionary origin, their codon usage converges toward reduced competition with highly expressed genes and improved translation during stress.

The model proposed here complements rather than replaces existing explanations for virulence-gene A/T richness. Horizontal acquisition, donor-genome composition and H-NS-mediated silencing all likely contribute to the composition of pathogenicity islands and other virulence loci. Our data unravel an additional layer: once acquired, virulence codon usage may be retained or evolve to support stress-adapted translation and to limit competition with highly expressed housekeeping genes.

More broadly, this work places synonymous codon usage at the interface between genome evolution, translation regulation and pathogenesis. It also identifies that specific virulence decoding strategies may expose vulnerabilities that are otherwise invisible under nutrient-rich laboratory conditions, suggesting tRNA charging and tRNA modifications as promising targets for broad anti-virulence strategies.

Future work using charged-tRNA profiling, ribosome profiling and chromosomal recoding of endogenous virulence loci will be needed to quantify the contribution of individual codons and tRNA isoacceptors to virulence-gene translation during infection. The present study establishes the genomic, experimental and in vivo basis for this model and provides a framework for testing how synonymous codon usage intersects with translation regulation and bacterial pathogenicity.

## Data availability

CodonPipe and associated analysis workflows are available at the Zenodo repositories cited in Methods. Supplementary Data are available at NAR Online.

- Data S1. Functional clusters per species
- Data S2. Sequences of recoded fluorescent protein genes
- Data S3. *S. enterica* SL1344 codon usage tables
- Data S4. Virulence clusters gene descriptions
- Data S5. pSLT and F plasmids subclusters gene lists
- Data S6. Fluorescence and OD measurements of recoded fluorescent proteins

## Acknowledgements

We thank Heng Wang for carrying out mice infection experiments and thank L. Van Melderen for critical reading of the manuscript.

## Author Contributions Statement

Conceptualization: FG, DB; Methodology: FG; Investigation: FG, DB; Visualization: FG; Funding acquisition: FG, DB; Project administration: FG, DB; Supervision: FG, DB; Writing – original draft: FG; Writing – review & editing: FG, DB

