## Supplementary data for "Stress-adapted codon usage enables efficient virulence-gene expression in *Salmonella*"

### Supplementary Figures

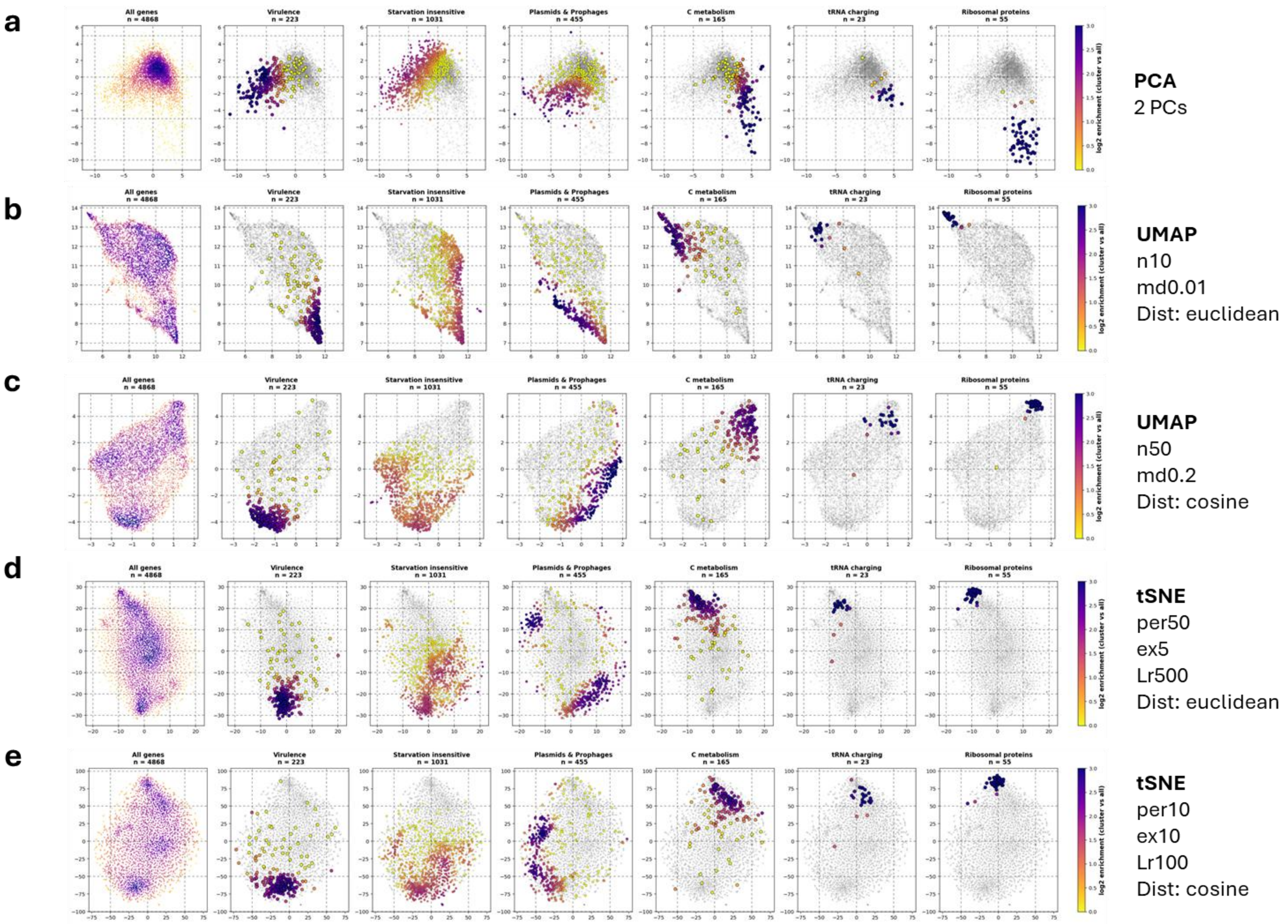
a
b
c
d
e
All genes
n = 4868
Virulence
n = 223
Starvation insensitive
n = 1031
Plasmids & Prophages
n = 455
C metabolism
n = 165
tRNA charging
n = 23
Ribosomal proteins
n = 55
log2 enrichment (cluster vs all)
PCA
2 PCs
UMAP
n10
md0.01
Dist: euclidean
UMAP
n50
md0.2
Dist: cosine
tSNE
per50
ex5
Lr500
Dist: euclidean
tSNE
per10
ex10
Lr100
Dist: cosine

**Figure S1. Codon-usage clustering is robust to dimensionality-reduction method and parameter choice.** Two-dimensional representations of *STm* coding sequences in the PCA (a), UMAP (b,c), and t-SNE (d,e) spaces with different parameter settings. All genes are shown together with representative functional gene clusters. All coding sequences are displayed in grey and representative functional gene clusters are represented by density enrichment plots. Color intensity indicates local enrichment relative to the whole-genome background.

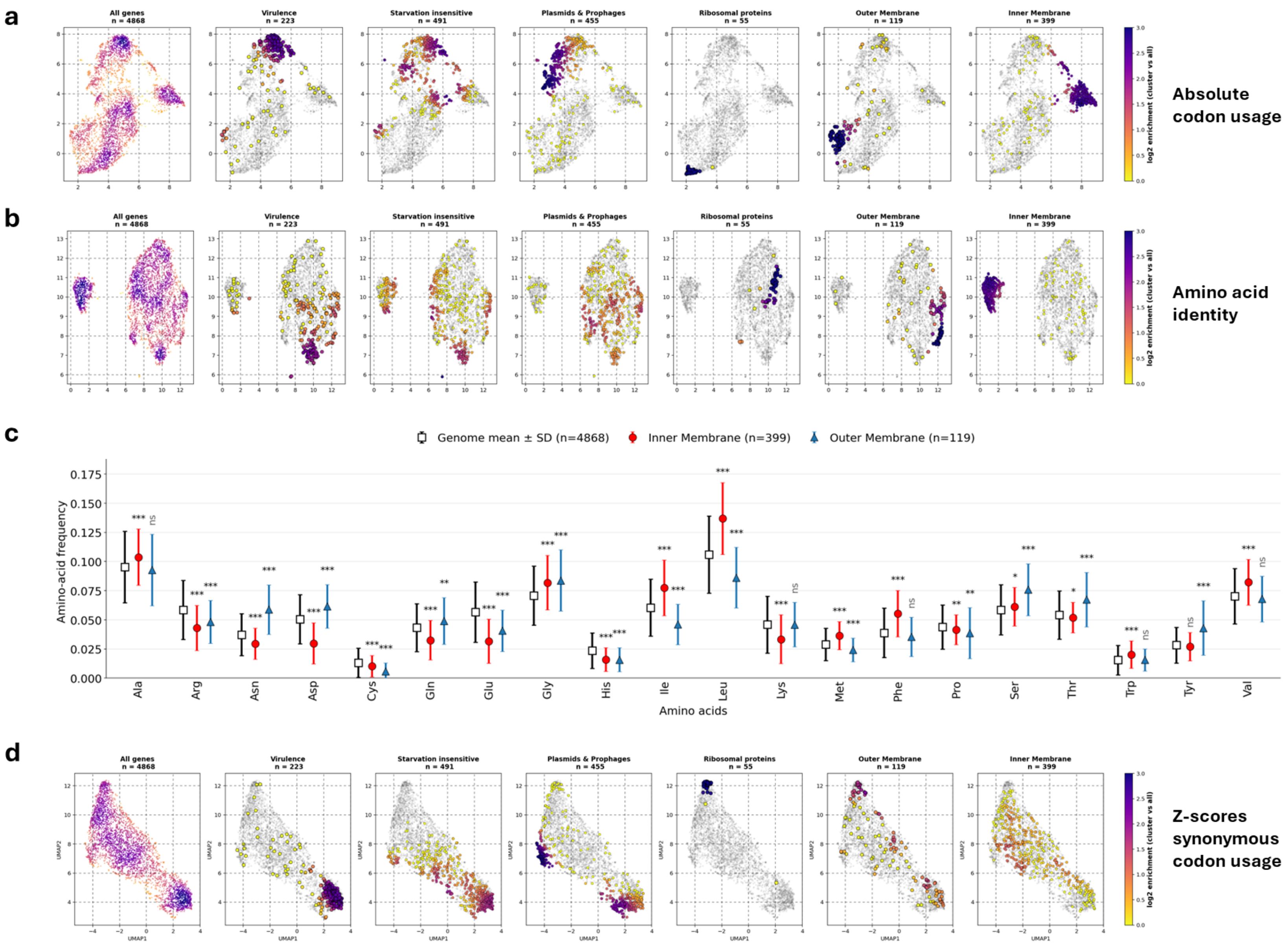
a
All genes n = 4868
Virulence n = 223
Starvation insensitive n = 491
Plasmids & Prophages n = 455
Ribosomal proteins n = 55
Outer Membrane n = 119
Inner Membrane n = 399
log2 enrichment (cluster vs all)
Absolute codon usage
b
All genes n = 4868
Virulence n = 223
Starvation insensitive n = 491
Plasmids & Prophages n = 455
Ribosomal proteins n = 55
Outer Membrane n = 119
Inner Membrane n = 399
log2 enrichment (cluster vs all)
Amino acid identity
c
Genome mean ± SD (n=4868)
Inner Membrane (n=399)
Outer Membrane (n=119)
Amino-acid frequency
Amino acids
Ala
Arg
Asn
Asp
Cys
Gln
Glu
Gly
His
Ile
Leu
Lys
Met
Phe
Pro
Ser
Thr
Trp
Tyr
Val
d
All genes n = 4868
Virulence n = 223
Starvation insensitive n = 491
Plasmids & Prophages n = 455
Ribosomal proteins n = 55
Outer Membrane n = 119
Inner Membrane n = 399
UMAP1
UMAP2
log2 enrichment (cluster vs all)
Z-scores synonymous codon usage

**Figure S2. Clustering on synonymous codon usage z-scores improves specificity and sensitivity.** Comparison of codon-usage clustering obtained using three codon usage metrics in *STm*: (a) absolute codon usage, (b) amino acid identity, and (d) genome-centered z-scores of synonymous codon usage. Functional overlays are shown for representative *Salmonella* gene clusters. In (a), (b) and (d), plots show all coding sequences in grey and density enrichment for representative functional gene clusters. Color intensity indicates local enrichment relative to the whole-genome background. Gene numbers are indicated above each panel. (c) Amino-acid composition of inner- and outer-membrane proteins compared with the genome-wide distribution. Data are average amino acid usage values and error bars are standard deviations. Statistical significance was assessed using two-tailed Student's *t*-tests: * $P < 0.05$, ** $P < 0.01$, *** $P < 0.001$.

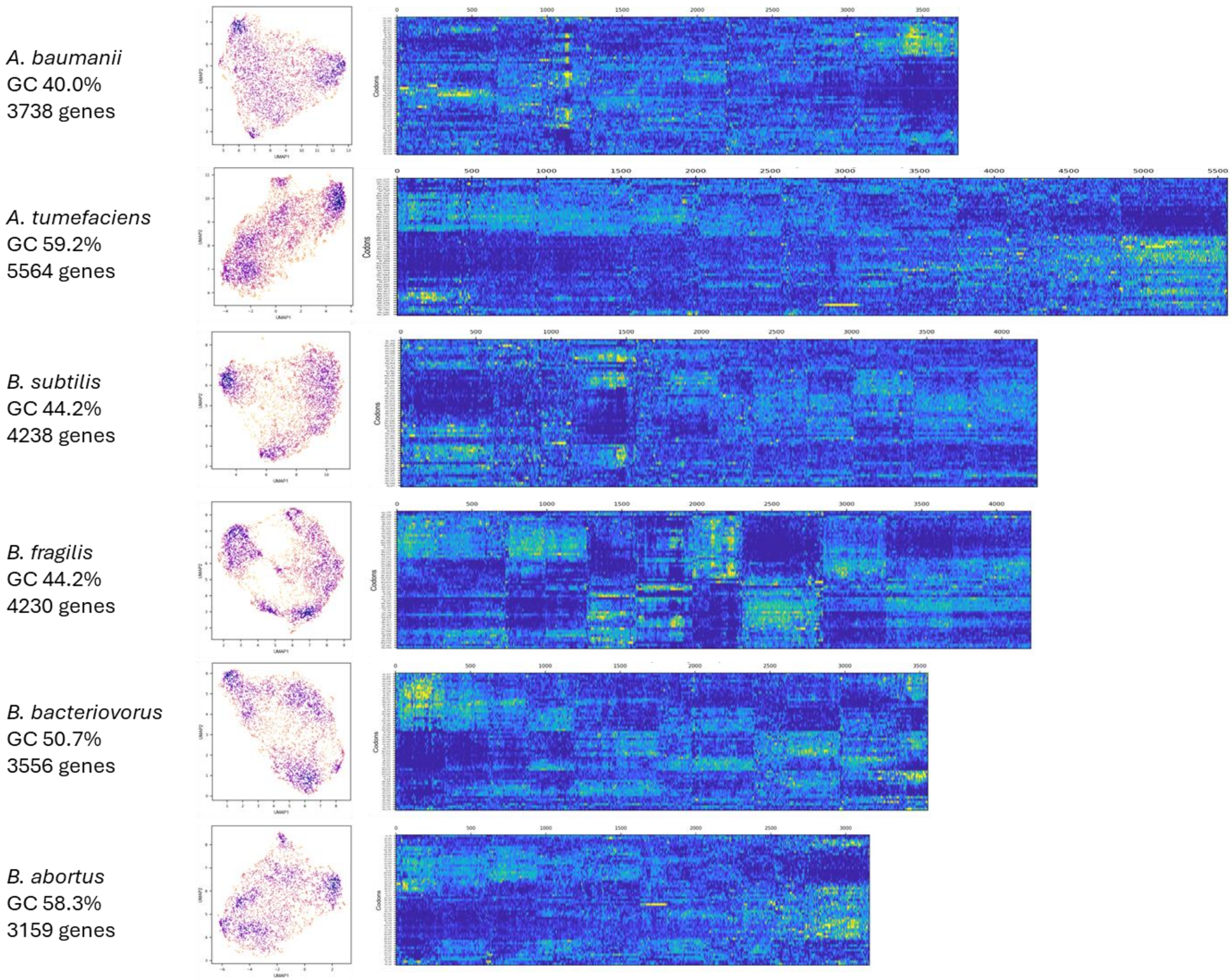
*A. baumanii*
GC 40.0%
3738 genes
*A. tumefaciens*
GC 59.2%
5564 genes
*B. subtilis*
GC 44.2%
4238 genes
*B. fragilis*
GC 44.2%
4230 genes
*B. bacteriovorus*
GC 50.7%
3556 genes
*B. abortus*
GC 58.3%
3159 genes

*B. pseudomallei*
GC 68.5%
6215 genes

*C. crescentus*
GC 67.7%
3879 genes

*C. indologenes*
GC 38.3%
4392 genes

*E. cloaceae*
GC 56.0%
4718 genes

*E. coli O157:H7*
GC 51.7%
5155 genes

*F. tullarensis*
GC 33.1%
1761 genes

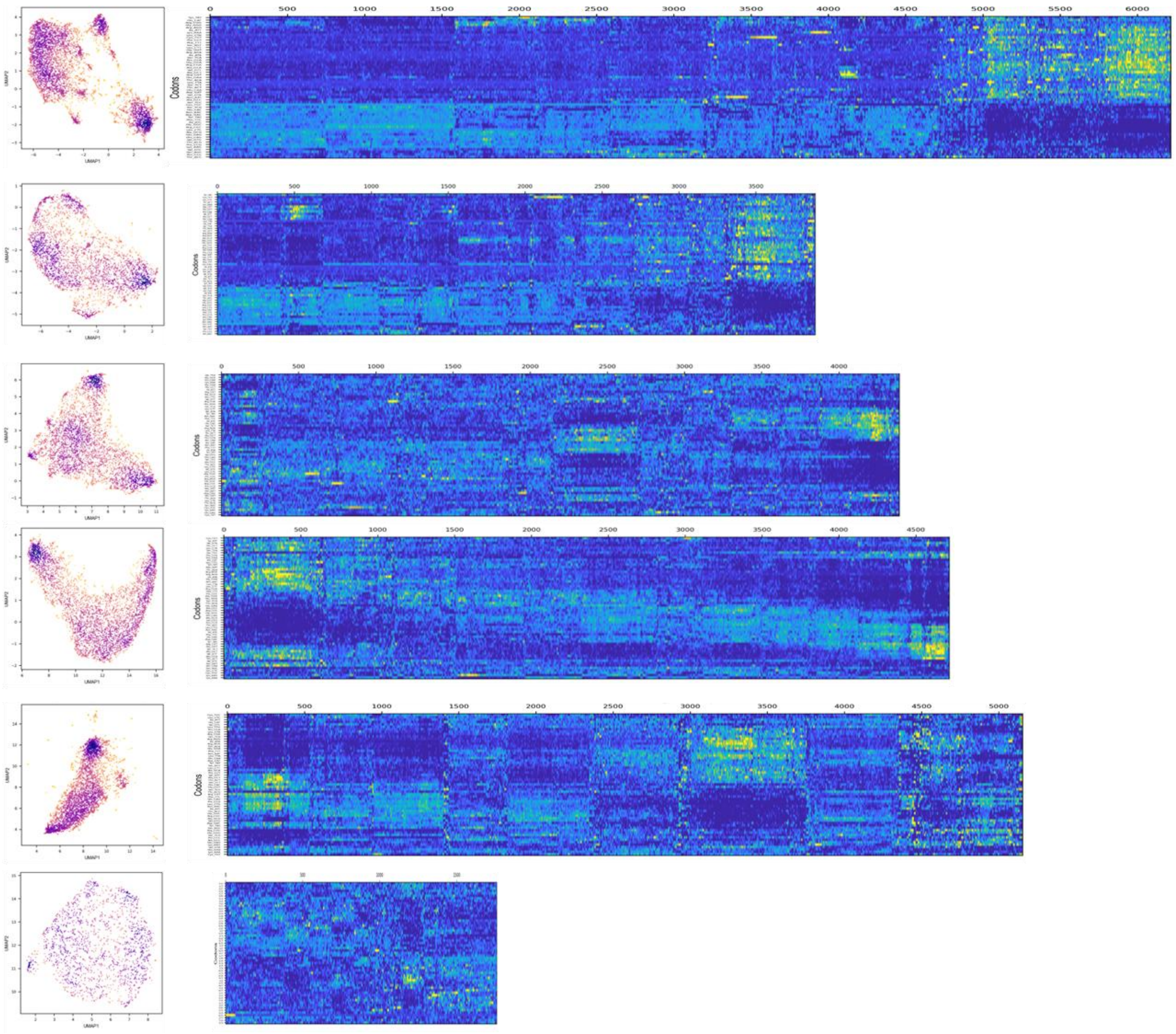

*H. pylori*
GC 39.2%
1584 genes

*K. pneumoniae*
GC 58.3%
5779 genes

*L. monocytogenes*
GC 38.4%
2867 genes

*M. tuberculosis*
GC 65.9%
3906 genes

*N. gonorrhoeae*
GC 53.8%
2207 genes

*N. meningitidis*
GC 53.3%
2143 genes

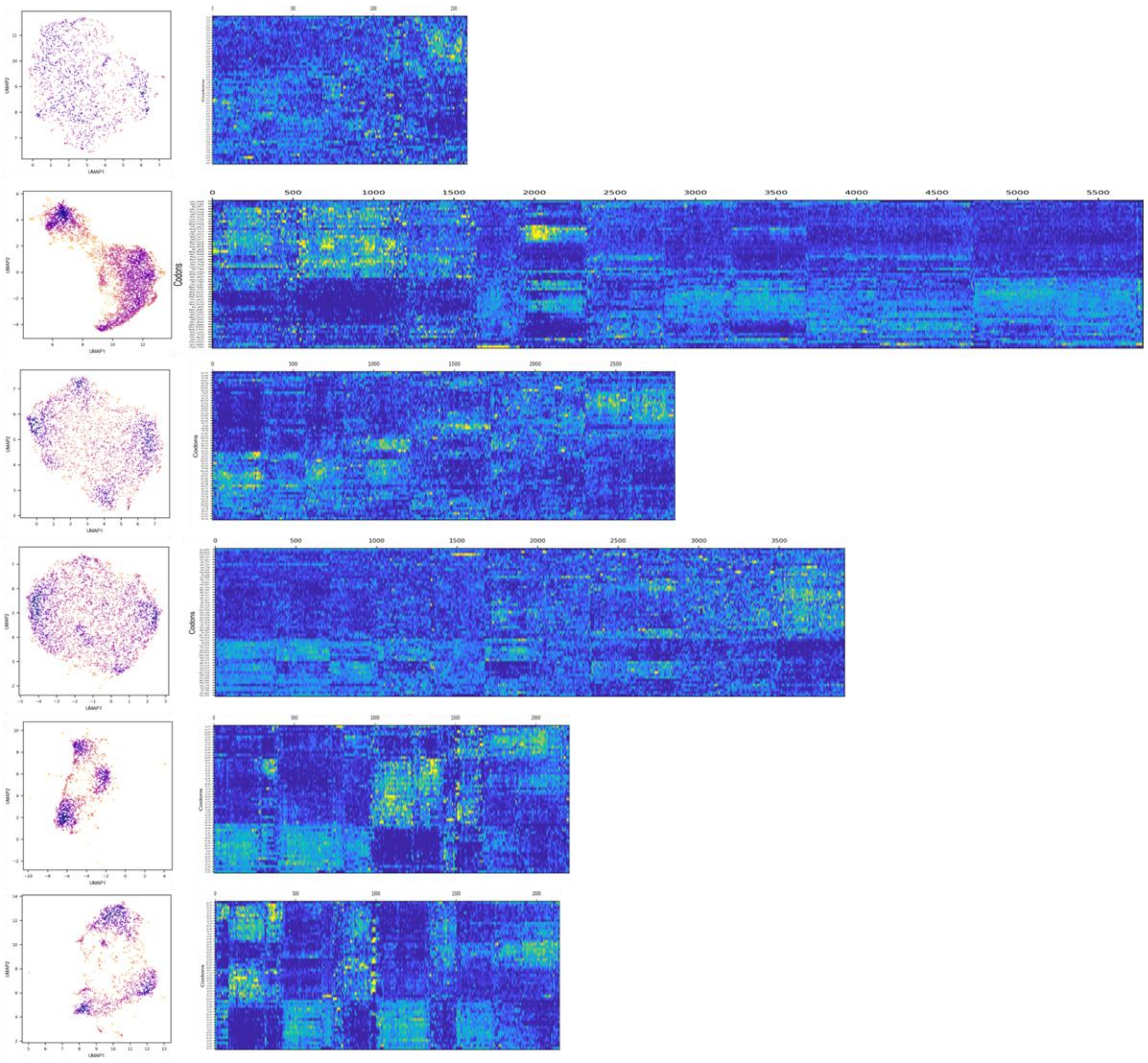

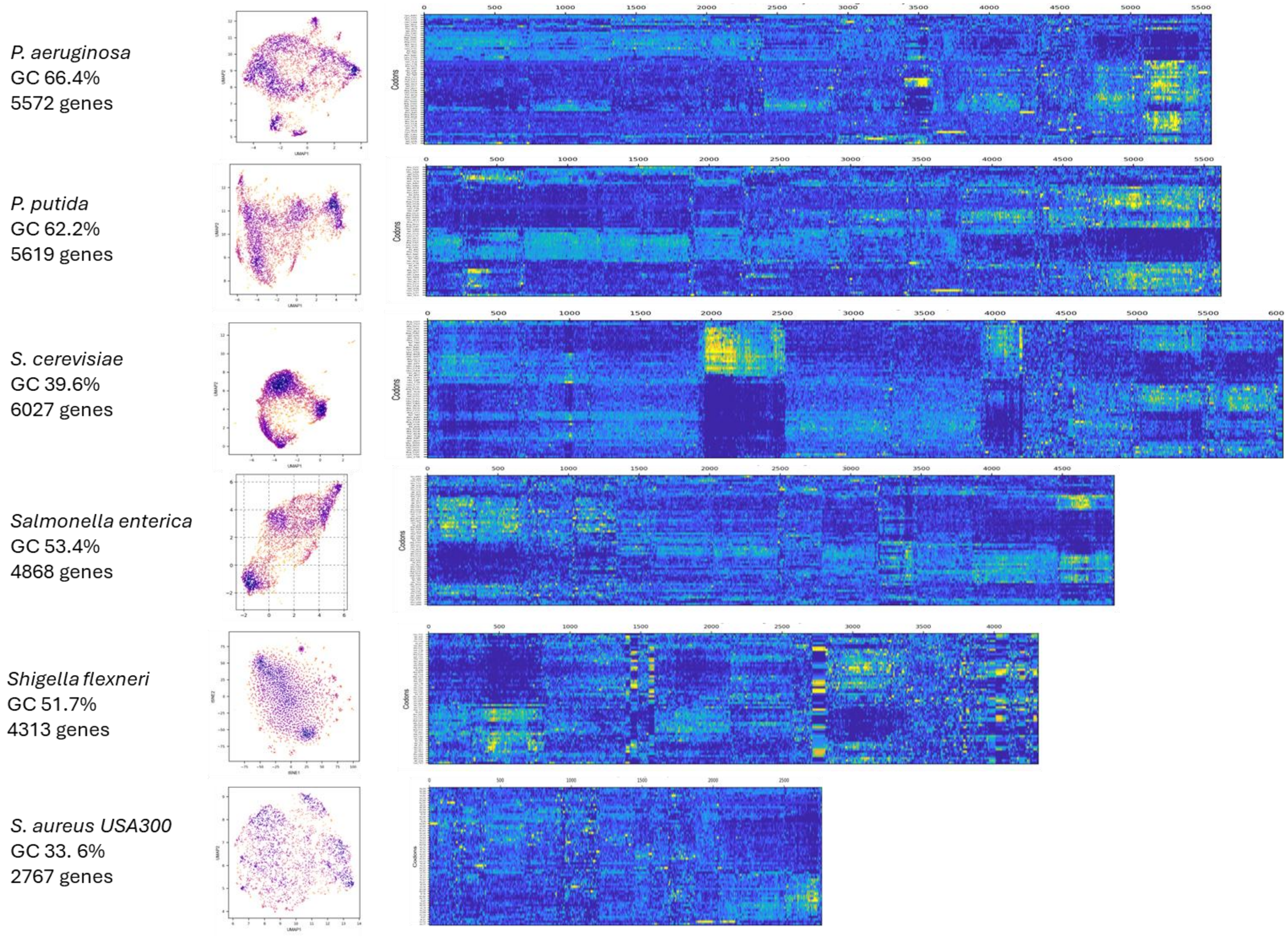
*P. aeruginosa*
GC 66.4%
5572 genes
*P. putida*
GC 62.2%
5619 genes
*S. cerevisiae*
GC 39.6%
6027 genes
*Salmonella enterica*
GC 53.4%
4868 genes
*Shigella flexneri*
GC 51.7%
4313 genes
*S. aureus USA300*
GC 33. 6%
2767 genes
Codons

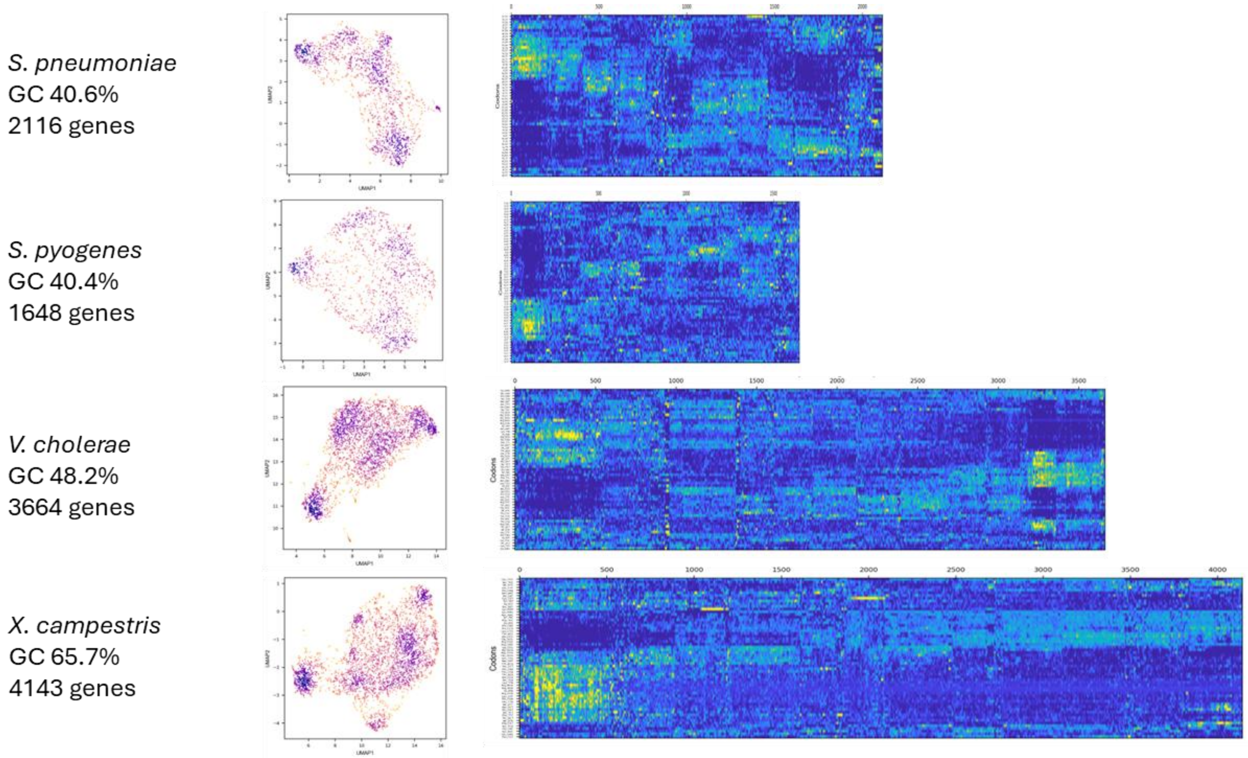


**Figure S3. Codon clustering reveals diverse and strong structures across bacterial genomes,** Overview of codon-usage landscapes across 28 bacterial species and *Saccharomyces cerevisiae*. For each organism, all genes were plotted in the 2D UMAP space and a heatmap of synonymous codon usage z-scores (ZCU) was plotted in the UMAP-reordered genome. Genome size and GC content are indicated for each species. Heatmap width is proportional to the genome size (in genes). Species are ordered alphabetically.

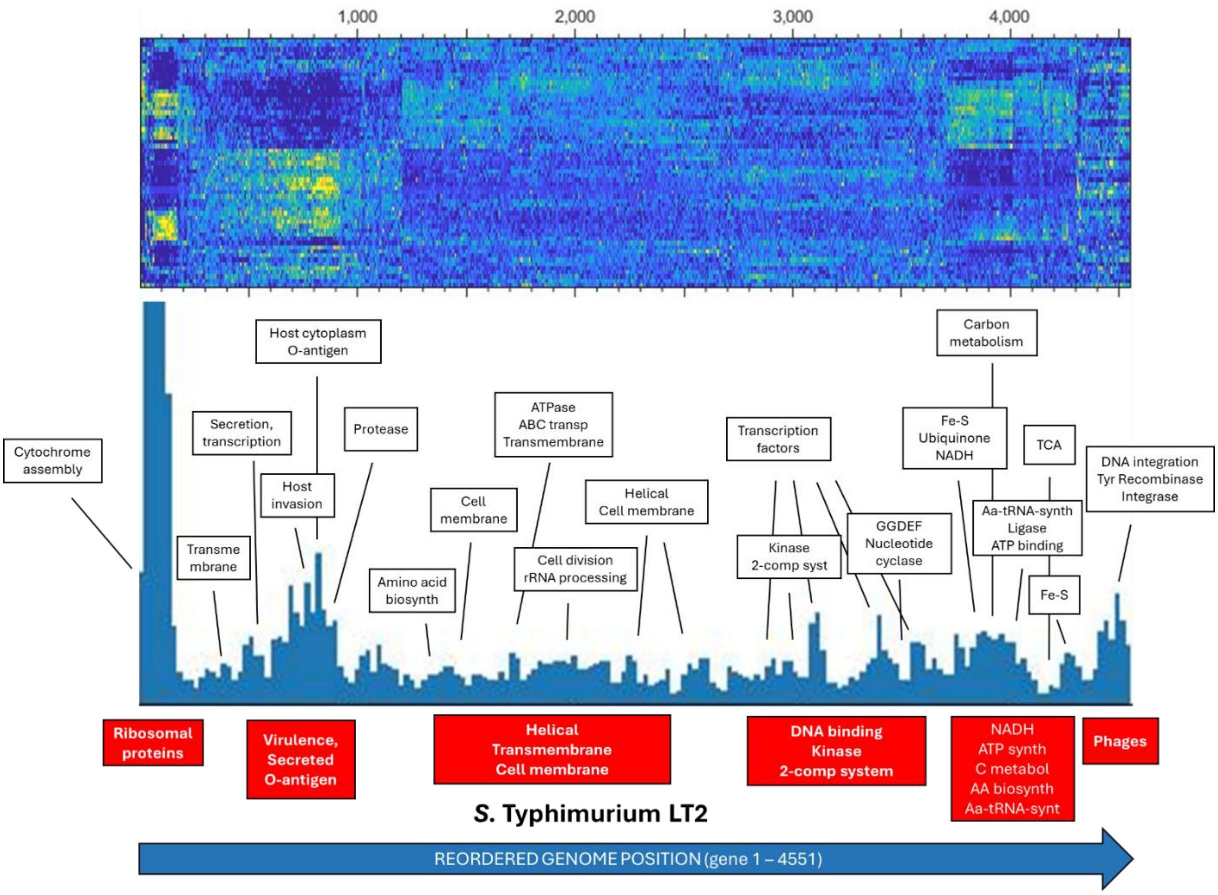


**Figure S4. Sliding window enrichment analysis in *Salmonella enterica* Typhimurium LT2.** *Salmonella* LT2 UMAP-reordered genome was scanned for functional enrichment using a sliding window of 100 genes advancing in steps of 25 genes across the codon-usage landscape. For each window, functional enrichment was assessed using DAVID (1), and enrichment peaks were plotted along the reordered genome axis. Major peaks were manually annotated according to enriched functional terms and genomic context. Functional enrichment peaks along the reordered genome axis are shown in blue and were manually annotated. Note that *Salmonella* LT2 was used in this example rather than SL1344 because SL1344 is not available in DAVID bioinformatics.

| *Salmonella* SL1344 pSLT cluster 2 | | | | |
|---|---|---|---|---|
| **Locus Tag** | **Gene** | **umap-x** | **umap-y** | **Protein description** |
| SL1344_RS23990 | | -2.12 | -0.85 | lysozyme |
| SL1344_RS24180 | repC | -2.12 | -1.07 | replication regulatory protein RepA |
| SL1344_RS23960 | | -2.09 | -0.96 | carbonic anhydrase |
| SL1344_RS23795 | traL | -1.94 | -0.91 | type IV conjugative transfer system protein TraL |
| SL1344_RS23675 | traS | -1.89 | -1.05 | entry exclusion protein |
| SL1344_RS23725 | | -1.87 | -0.91 | hypothetical protein |
| SL1344_RS24075 | | -1.79 | -1.45 | hypothetical protein |
| SL1344_RS27530 | | -1.69 | -1.16 | DUF932 domain-containing protein |
| SL1344_RS27815 | | -1.58 | -1.52 | hypothetical protein |
| SL1344_RS24010 | spvD | -1.57 | -1.68 | SPI-2 type III secretion system effector cysteine hydrolase SpvD |
| SL1344_RS27935 | | -1.54 | -2.17 | replication protein RepA |
| SL1344_RS23805 | traJ | -1.53 | -1.29 | PAS domain-containing protein |
| SL1344_RS23920 | parB | -1.51 | -1.39 | ParB family protein |
| SL1344_RS24170 | Tap | -1.41 | -2.14 | RepA leader peptide Tap |
| SL1344_RS23810 | traM | -1.37 | -1.58 | conjugal transfer relaxosome DNA-binding protein TraM |
| SL1344_RS23770 | trbD | -1.07 | -1.23 | DUF2689 domain-containing protein |
| SL1344_RS23985 | spvR | -0.92 | -1.59 | virulence genes transcriptional activator SpvR |
| SL1344_RS26895 | | -0.89 | -1.62 | hypothetical protein |
| SL1344_RS27315 | | -0.86 | -1.13 | site-specific integrase |
| SL1344_RS24125 | orf6 | -0.85 | -1.21 | hypothetical protein |
| SL1344_RS24005 | spvC | -0.83 | -0.96 | type III secretion system effector phosphothreonine lyase SpvC |
| SL1344_RS24000 | spvB | -0.83 | -1.72 | SPI-2 type III secretion system effector NAD(+)--protein-arginine ADP-ribosyltransferase SpvB |
| SL1344_RS24100 | pefB | -0.79 | -1.71 | transcriptional regulator PefB |
| SL1344_RS23815 | mltE | -0.74 | -1.69 | transglycosylase SLT domain-containing protein MltE |
| SL1344_RS23860 | | -0.47 | -1.25 | DNA polymerase III subunit theta |
| SL1344_RS26285 | traY | -0.41 | -1.16 | conjugal transfer relaxosome protein TraY |
| SL1344_RS27885 | | -0.3 | -1.35 | hypothetical protein |
| SL1344_RS26320 | tnpA | -0.28 | -0.96 | IS200/IS605 family transposase |
| SL1344_RS23980 | | -0.22 | -1.58 | IS630 family transposase |
| SL1344_RS24095 | | -0.13 | -1.28 | NirD/YgiW/YdeI family stress tolerance protein |
| SL1344_RS24155 | srgC | -0.06 | -1.15 | AraC family transcriptional regulator |
| SL1344_RS23925 | parA | -0.02 | -1.54 | AAA family ATPase |
| SL1344_RS23695 | traQ | 0.25 | -0.26 | type-F conjugative transfer system pilin chaperone TraQ |
| SL1344_RS23965 | rlgA | 0.25 | -1.47 | recombinase family protein |
| SL1344_RS23705 | trbE | 0.45 | -1.48 | conjugal transfer protein TrbE |
| SL1344_RS27805 | | 0.46 | -1.27 | hypothetical protein |
| SL1344_RS24085 | repA2 | 0.54 | -1.14 | RepB family plasmid replication initiator protein |
| SL1344_RS23950 | tlpA | 0.63 | -1.34 | DNA-binding protein |
| SL1344_RS24185 | | 0.66 | -0.9 | phospholipase D family protein |
| SL1344_RS23670 | traT | 0.97 | -0.4 | conjugal transfer complement resistance protein TraT |

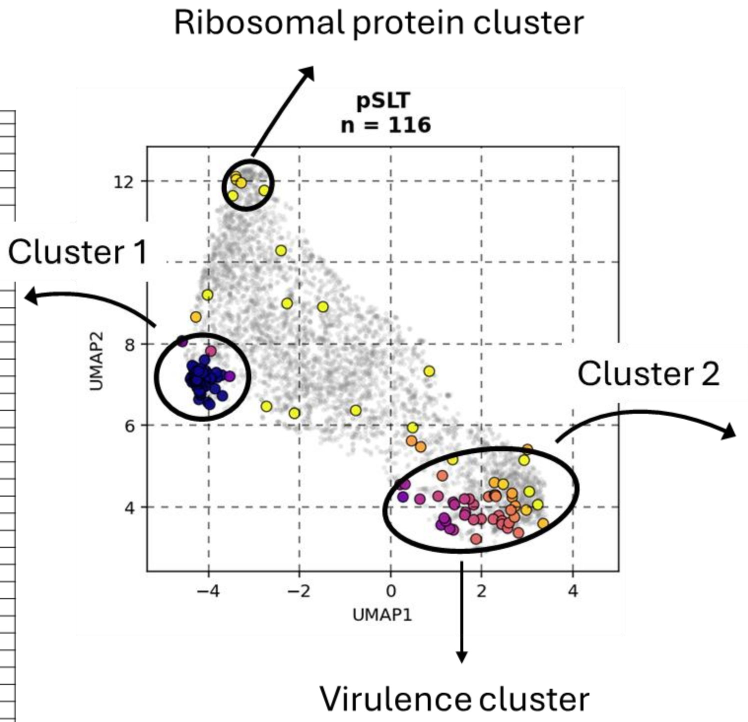


| *Salmonella* SL1344 pSLT cluster 1 | | | | |
|---|---|---|---|---|
| **Locus Tag** | **Gene** | **umap-x** | **umap-y** | **Protein description** |
| SL1344_RS24120 | orf5 | 1.97 | -0.21 | membrane protein |
| SL1344_RS23975 | | 2.23 | -0.05 | ExeA family protein |
| SL1344_RS23850 | ssbB | 2.25 | 0.72 | single-stranded DNA-binding protein SSB2 |
| SL1344_RS23750 | | 2.25 | 0.06 | hypothetical protein |
| SL1344_RS23880 | | 2.32 | 0.43 | DUF1380 family protein |
| SL1344_RS23760 | traR | 2.46 | 0.32 | conjugal transfer protein TraR |
| SL1344_RS23935 | | 2.57 | 0.29 | DUF2913 family protein |
| SL1344_RS23775 | traP | 2.61 | 0.3 | conjugal transfer pilus-stabilizing protein TraP |
| SL1344_RS23945 | | 2.62 | 0.11 | 3'-5' exonuclease |
| SL1344_RS23915 | samA | 2.64 | 0.11 | Y-family DNA polymerase |
| SL1344_RS23690 | trbB | 2.68 | 0.15 | type-F conjugative transfer system pilin assembly thiol-disulfide isomerase TrbB |
| SL1344_RS24160 | | 2.74 | 0.53 | SdiA-regulated domain-containing protein |
| SL1344_RS23680 | traG | 2.76 | 0.33 | conjugal transfer mating-pair stabilization protein TraG |
| SL1344_RS23910 | umuD | 2.77 | 0.19 | translesion error-prone DNA polymerase V autoproteolytic subunit |
| SL1344_RS23970 | rlgA | 2.79 | 0.42 | IS481 family transposase |
| SL1344_RS23955 | | 2.8 | 0.23 | hypothetical protein |
| SL1344_RS23720 | trbC | 2.91 | 0.42 | type-F conjugative transfer system pilin assembly protein TrbC |
| SL1344_RS23755 | | 2.94 | 0.6 | hypothetical protein |
| SL1344_RS24145 | srgB | 2.96 | 0.58 | SdiA-regulated domain-containing protein |
| SL1344_RS24055 | | 2.96 | 0.64 | hypothetical protein |
| SL1344_RS23660 | vapC | 2.98 | 0.48 | type II toxin-antitoxin system tRNA(fMet)-specific endonuclease VapC |
| SL1344_RS24045 | | 2.98 | 1.05 | EAL domain-containing protein |
| SL1344_RS24150 | rck | 3.01 | 0.4 | complement resistance protein Rck |
| SL1344_RS24050 | rsdB | 3.02 | 0.58 | site-specific integrase |
| SL1344_RS23665 | traD | 3.03 | 0.56 | type IV conjugative transfer system coupling protein TraD |
| SL1344_RS24165 | repA | 3.04 | 0.43 | incFII family plasmid replication initiator RepA |
| SL1344_RS23835 | psiB | 3.08 | 0.63 | conjugation system SOS inhibitor PsiB |
| SL1344_RS24070 | ccdA | 3.09 | 0.58 | type II toxin-antitoxin system antitoxin CcdA |
| SL1344_RS23710 | traN | 3.09 | 0.54 | type-F conjugative transfer system mating-pair stabilization protein TraN |
| SL1344_RS23780 | traB | 3.09 | 0.51 | F-type conjugal transfer pilus assembly protein TraB |
| SL1344_RS24140 | srgA | 3.12 | 0.61 | DsbA family protein |
| SL1344_RS23740 | trbI | 3.12 | 0.77 | type-F conjugative transfer system protein TrbI |
| SL1344_RS23875 | | 3.15 | 0.63 | membrane protein |
| SL1344_RS23765 | traV | 3.15 | 0.63 | type IV conjugative transfer system lipoprotein TraV |
| SL1344_RS23685 | traH | 3.16 | 0.7 | conjugal transfer pilus assembly protein TraH |
| SL1344_RS24135 | orf7 | 3.17 | 0.77 | helix-turn-helix domain-containing protein |
| SL1344_RS23650 | traI | 3.17 | 0.62 | conjugative transfer relaxase/helicase TraI |
| SL1344_RS23785 | traK | 3.22 | 0.63 | type-F conjugative transfer system secretin TraK |
| SL1344_RS23840 | | 3.22 | 0.8 | ParB/RepB/Spo0J family partition protein |
| SL1344_RS24035 | | 3.23 | 0.84 | membrane protein |
| SL1344_RS24040 | | 3.23 | 0.81 | CaiF/GrlA family transcriptional regulator |
| SL1344_RS23745 | traC | 3.25 | 0.74 | type IV secretion system protein TraC |
| SL1344_RS24065 | ccdB | 3.26 | 0.81 | type II toxin-antitoxin system toxin CcdB |
| SL1344_RS24115 | pefD | 3.28 | 0.7 | fimbrial chaperone PefD |
| SL1344_RS23735 | traW | 3.29 | 0.85 | type-F conjugative transfer system protein TraW |
| SL1344_RS24110 | pefC | 3.29 | 0.83 | outer membrane usher protein PefC |
| SL1344_RS23700 | traF | 3.29 | 0.86 | type-F conjugative transfer system pilin assembly protein TraF |
| SL1344_RS23730 | traU | 3.31 | 0.82 | conjugal transfer pilus assembly protein TraU |
| SL1344_RS27810 | | 3.32 | 1.3 | LysM peptidoglycan-binding domain-containing protein |
| SL1344_RS23645 | traX | 3.32 | 0.84 | conjugal transfer pilus acetylase TraX |
| SL1344_RS23995 | spvA | 3.36 | 1.4 | virulence protein SpvA |
| SL1344_RS23830 | psiA | 3.37 | 0.88 | plasmid SOS inhibition protein A |
| SL1344_RS24060 | | 3.37 | 1.07 | hypothetical protein |
| SL1344_RS23640 | finO | 3.37 | 0.87 | fertility inhibition protein FinO |
| SL1344_RS24190 | | 3.43 | 1.16 | hypothetical protein |
| SL1344_RS23930 | samB | 3.44 | 1.16 | Rpn family recombination-promoting nuclease/putative transposase |
| SL1344_RS23715 | | 3.58 | 1.51 | HNH endonuclease |

**Figure S5. The *Salmonella* virulence plasmid pSLT separates into two codon-usage subclusters.** Genes encoded by the *Salmonella* pSLT plasmid were mapped onto the two-dimensional codon-usage UMAP space. pSLT genes did not form a single homogeneous group but separated into two distinct subclusters. Clusters compositions are provided in Data S5.

| Locus Tag | Gene | Umap-x | Umap-y | Protein description |
|---|---|---|---|---|
| AA953_RS23375 | trbJ | 13.86 | 1.06 | P-type conjugative transfer protein TrbJ |
| AA953_RS23510 | ccdA | 13.77 | 0.99 | type II toxin-antitoxin system antitoxin CcdA |
| AA953_RS23270 | traL | 14.02 | 0.90 | type IV conjugative transfer system protein TraL |
| AA953_RS23415 | vapB | 13.20 | 0.57 | toxin-antitoxin system antitoxin VapB |
| AA953_RS23250 | traM | 14.44 | 0.46 | conjugaltransfer relaxosome DNA-binding protein TraM |
| AA953_RS23380 | trbF | 14.53 | 0.43 | F-type conjugal transfer protein TrbF |
| AA953_RS23240 | | 14.87 | 0.34 | DUF932 domain-containing protein |
| AA953_RS23460 | tap | 14.35 | 0.34 | RepA leader peptide Tap |
| AA953_RS23495 | | 14.27 | 0.29 | TIR domain-containing protein |
| AA953_RS23535 | | 14.78 | 0.29 | ParB/RepB/Spo0J family plasmid partition protein |
| AA953_RS23530 | sopA | 14.66 | 0.22 | plasmid-partitioning protein SopA |
| AA953_RS26905 | | 13.89 | 0.07 | hypothetical protein |
| AA953_RS23295 | | 14.99 | 0.02 | conjugaltransfer protein TrbD |
| AA953_RS23360 | | 14.64 | 0.00 | protein ArtA |
| AA953_RS27395 | | 15.57 | -0.01 | hypothetical protein |
| AA953_RS26890 | | 14.19 | -0.01 | 3'-5' exonuclease |
| AA953_RS23290 | traP | 14.13 | -0.03 | conjugaltransfer pilus-stabilizing protein TraP |
| AA953_RS23500 | | 14.08 | -0.03 | KAP family P-loop NTPase fold protein |
| AA953_RS23485 | ppl | 13.42 | -0.04 | anti-phage protein Ppl |
| AA953_RS23520 | | 13.20 | -0.06 | site-specific integrase |
| AA953_RS26180 | | 13.30 | -0.06 | hypothetical protein |
| AA953_RS23260 | traY | 14.78 | -0.09 | conjugaltransfer relaxosome DNA-bindin protein TraY |
| AA953_RS23370 | trbB | 13.43 | -0.12 | type-F conjugative transfer system pilin assembly thiol-disulfide isomerase TrbB |
| AA953_RS27515 | | 13.74 | -0.14 | single-stranded DNA-binding protein |
| AA953_RS23450 | | 13.49 | -0.18 | Hok/Gef family protein |
| AA953_RS27510 | | 13.56 | -0.21 | hypothetical protein |
| AA953_RS23255 | traJ | 14.43 | -0.22 | conjugaltransfer transcriptional regulator TraJ |
| AA953_RS23445 | | 15.04 | -0.24 | hypothetical protein |
| AA953_RS23475 | | 14.91 | -0.34 | hypothetical protein |
| AA953_RS27200 | | 14.32 | -0.38 | DUF5431 family protein |
| AA953_RS23435 | | 12.88 | -0.39 | hypothetical protein |
| AA953_RS25595 | | 14.41 | -0.40 | recombinase family protein |
| AA953_RS23300 | trbG | 14.88 | -0.44 | conjugaltransfer protein TrbG |
| AA953_RS23525 | repE | 12.79 | -0.49 | replication initiation protein RepE |
| AA953_RS23620 | | 13.73 | -0.50 | type I toxin-antitoxin system Hok family toxin |
| AA953_RS23465 | | 14.29 | -0.54 | hypothetical protein |
| AA953_RS25575 | traS | 14.77 | -0.56 | conjugaltransfer entry exclusion protein TraS |
| AA953_RS23345 | | 14.06 | -0.59 | conjugaltransfer protein TrbE |
| AA953_RS26885 | | 14.66 | -0.60 | thermonuclease family protein |
| AA953_RS23490 | | 14.86 | -0.67 | hypothetical protein |
| AA953_RS23455 | | 14.19 | -0.71 | replication regulatory protein RepA |
| AA953_RS23355 | trbA | 14.09 | -0.79 | conjugaltransfer protein TrbA |

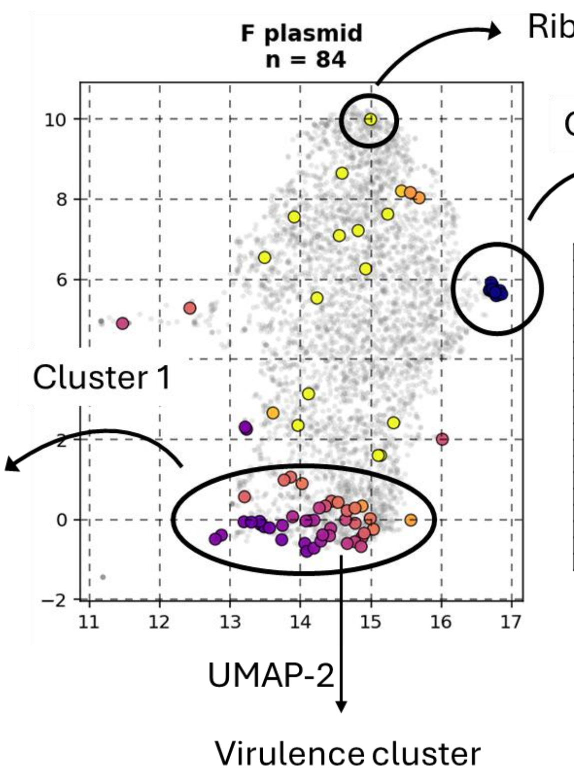


| Locus Tag | Gene | Umap-x | Umap-y | Protein description |
|---|---|---|---|---|
| AA953_RS23555 | | 16.71 | 5.92 | hypothetical protein |
| AA953_RS23470 | repA | 16.73 | 5.85 | incFII family plasmid replication initiator RepA |
| AA953_RS23405 | traD | 16.73 | 5.76 | type IV conjugative transfer system coupling protein TraD |
| AA953_RS23575 | | 16.73 | 5.75 | DUF1380 family protein |
| AA953_RS23325 | traW | 16.69 | 5.74 | type-F conjugative transfer system protein TraW |
| AA953_RS23285 | traB | 16.68 | 5.73 | F-type conjugal transfer pilus assembly protein TraB |
| AA953_RS23515 | ccdB | 16.83 | 5.72 | type II toxin-antitoxin system toxin CcdB |
| AA953_RS23505 | pifC | 16.73 | 5.72 | transcriptional repressor PifC |
| AA953_RS23315 | traC | 16.72 | 5.70 | type IV secretion system protein TraC |
| AA953_RS23420 | traI | 16.74 | 5.69 | conjugative transfer relaxase/helicase TraI |
| AA953_RS23610 | | 16.77 | 5.68 | plasmid SOS inhibition protein A |
| AA953_RS23340 | traN | 16.79 | 5.65 | type-F conjugative transfer system mating-pair stabilization protein TraN |
| AA953_RS23385 | traH | 16.76 | 5.65 | conjugaltransfer pilus assembly protein TraH |
| AA953_RS23605 | psiB | 16.86 | 5.63 | conjugation system SOS inhibitor PsiB |
| AA953_RS23425 | traX | 16.82 | 5.63 | conjugaltransfer pilus acetylase TraX |
| AA953_RS23390 | traG | 16.82 | 5.62 | conjugaltransfer mating-pair stabilization protein TraG |
| AA953_RS23335 | trbC | 16.81 | 5.62 | type-F conjugative transfer system pilin assembly protein TrbC |
| AA953_RS23350 | traF | 16.80 | 5.60 | type-F conjugative transfer system pilin assembly protein TraF |
| AA953_RS23430 | | 16.78 | 5.59 | fertility inhibition protein FinO |

**Figure S6. The *E. coli* F plasmid separates into cargo/accessory and maintenance/transfer codon-usage subclusters.** Genes from the *E. coli* F plasmid were projected onto the two-dimensional codon-usage UMAP space. Similar to the *Salmonella* pSLT plasmid, F-plasmid genes formed two main subclusters. Clusters compositions are provided in Data S5.

**a**

***Listeria monocytogenes***

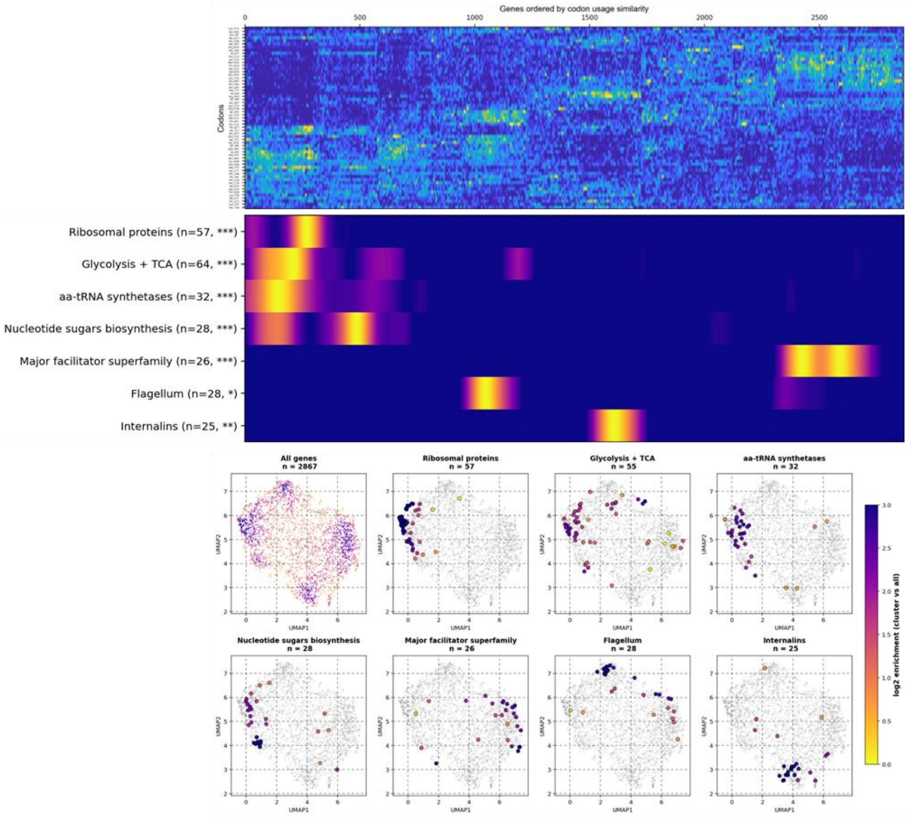


**b**

***Pseudomonas aeruginosa PAO1***

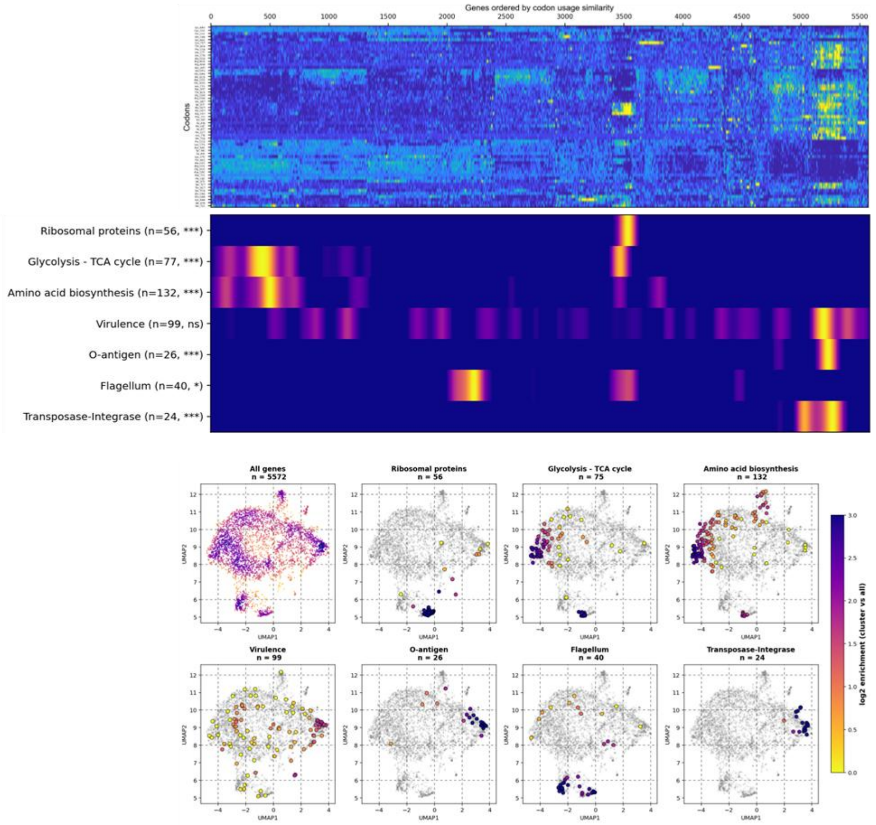

**c** ***Burkholderia pseudomallei***

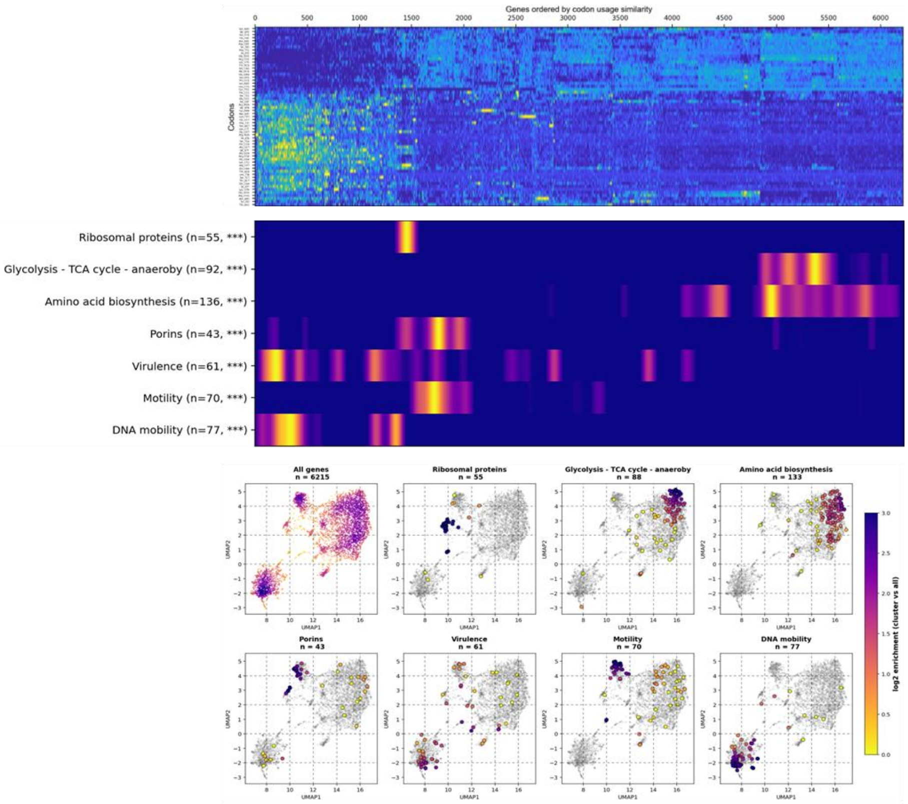


**d** ***Bacillus subtilis***

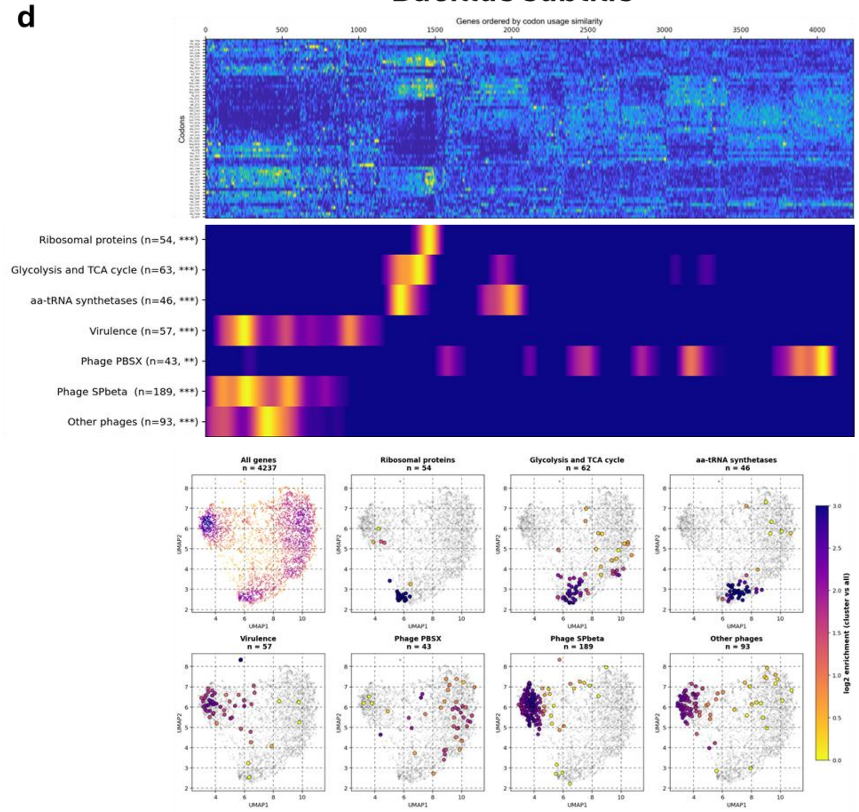

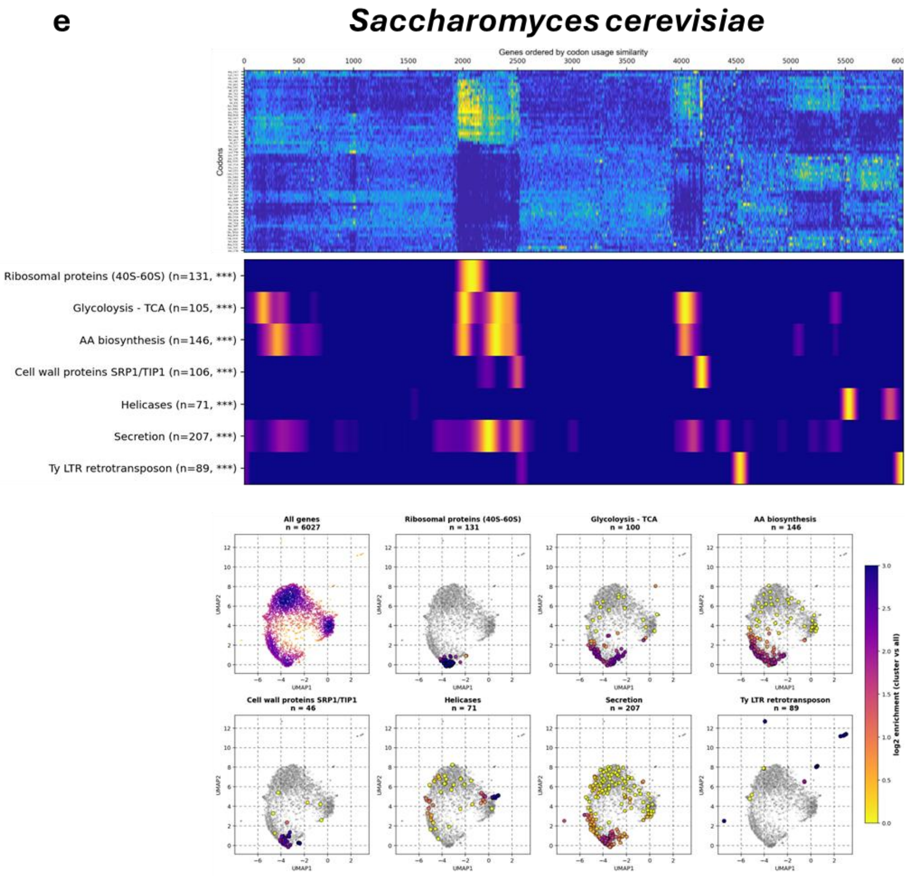


**Figure S7. Codon usage forms functional clusters across diverse bacterial and eukaryotic genomes.** CodonPipe analysis of *Listeria monocytogenes*, *Pseudomonas aeruginosa* PAO1, *Burkholderia pseudomallei*, *Bacillus subtilis* and *Saccharomyces cerevisiae*. For each organism, coding sequences were reordered according to synonymous codon-usage similarity, and functional gene sets were mapped both along the reordered genome axis and onto the two-dimensional UMAP space. Number of genes are indicated and statistical significance of non-random cluster localization was tested using a 1D-KS test (* $p<0.05$, ** $p<0.01$, *** $p<0.001$). Gene clusters compositions are provided in Data S1.

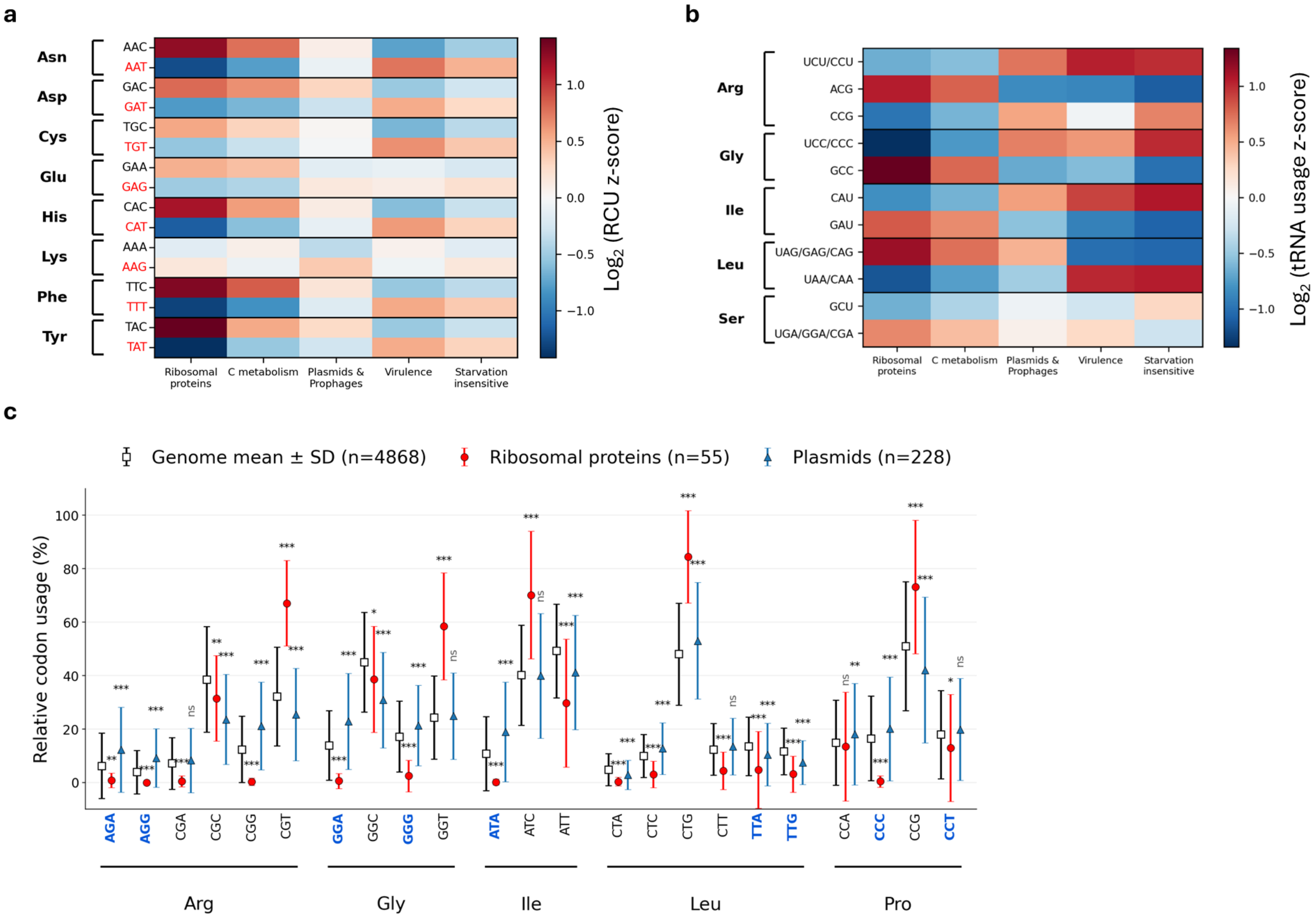
a
Asn
Asp
Cys
Glu
His
Lys
Phe
Tyr
AAC
AAT
GAC
GAT
TGC
TGT
GAA
GAG
CAC
CAT
AAA
AAG
TTC
TTT
TAC
TAT
Ribosomal proteins
C metabolism
Plasmids & Prophages
Virulence
Starvation insensitive
Log2 (RCU z-score)
1.0
0.5
0.0
−0.5
−1.0
b
Arg
Gly
Ile
Leu
Ser
UCU/CCU
ACG
CCG
UCC/CCC
GCC
CAU
GAU
UAG/GAG/CAG
UAA/CAA
GCU
UGA/GGA/CGA
Ribosomal proteins
C metabolism
Plasmids & Prophages
Virulence
Starvation insensitive
Log2 (tRNA usage z-score)
1.0
0.5
0.0
−0.5
−1.0
c
Genome mean ± SD (n=4868)
Ribosomal proteins (n=55)
Plasmids (n=228)
Relative codon usage (%)
100
80
60
40
20
0
AGA
AGG
CGA
CGC
CGG
CGT
GGA
GGC
GGG
GGT
ATA
ATC
ATT
CTA
CTC
CTG
CTT
TTA
TTG
CCA
CCC
CCG
CCT
Arg
Gly
Ile
Leu
Pro

**Figure S8. Decoding strategies differ across major *Salmonella* gene clusters.** Comparison of codon-decoding strategies across representative *STm* gene clusters. (a) Heatmap showing Watson–Crick versus wobble codon preferences for amino-acid families decoded by a single tRNA and containing two synonymous codons. Wobble-decoded codons are indicated in red. (b) Heatmap showing shifts in tRNA usage across major gene clusters for codon families decoded by multiple tRNA isoacceptors. When individual codons could not be unambiguously assigned to a single tRNA, tRNA isoacceptors were grouped into pooled decoding units. (c) Relative usage of starvation-insensitive codons in plasmid-associated genes compared with ribosomal genes and the genome-wide distribution. Starvation-insensitive codons are indicated in blue. Data are means ± SD. Statistical significance was assessed using two-tailed Student's *t*-tests. *$P < 0.05$, **$P < 0.01$, ***$P < 0.001$.

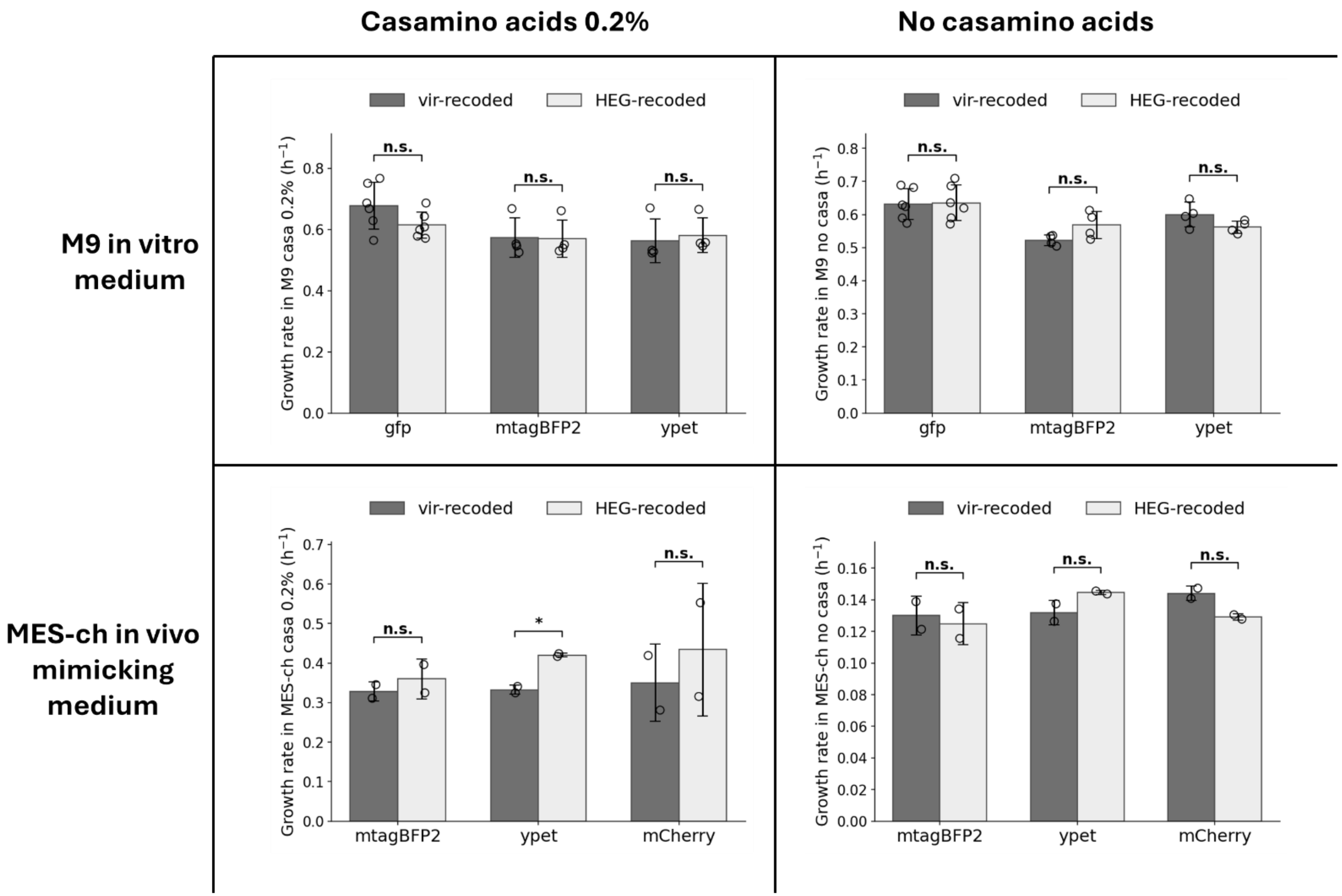


**Figure S9. Recoded fluorescent-protein reporters do not impair *STm* fitness in vitro.** Exponentially growing bacteria were diluted into 1 mL of the indicated medium in 24-well microplates, and $OD_{600}$ was monitored every 15 min for 18 h. Bar graphs show growth rates (in $h^{-1}$) of *STm* carrying vir- or HEG-recoded fluorescent protein reporters in M9 or MES-ch, with or without casamino acids 0.2%. Error bars are SD, and dots represent biological replicates (n). Statistical tests are unpaired two-tailed Student’s t-tests.

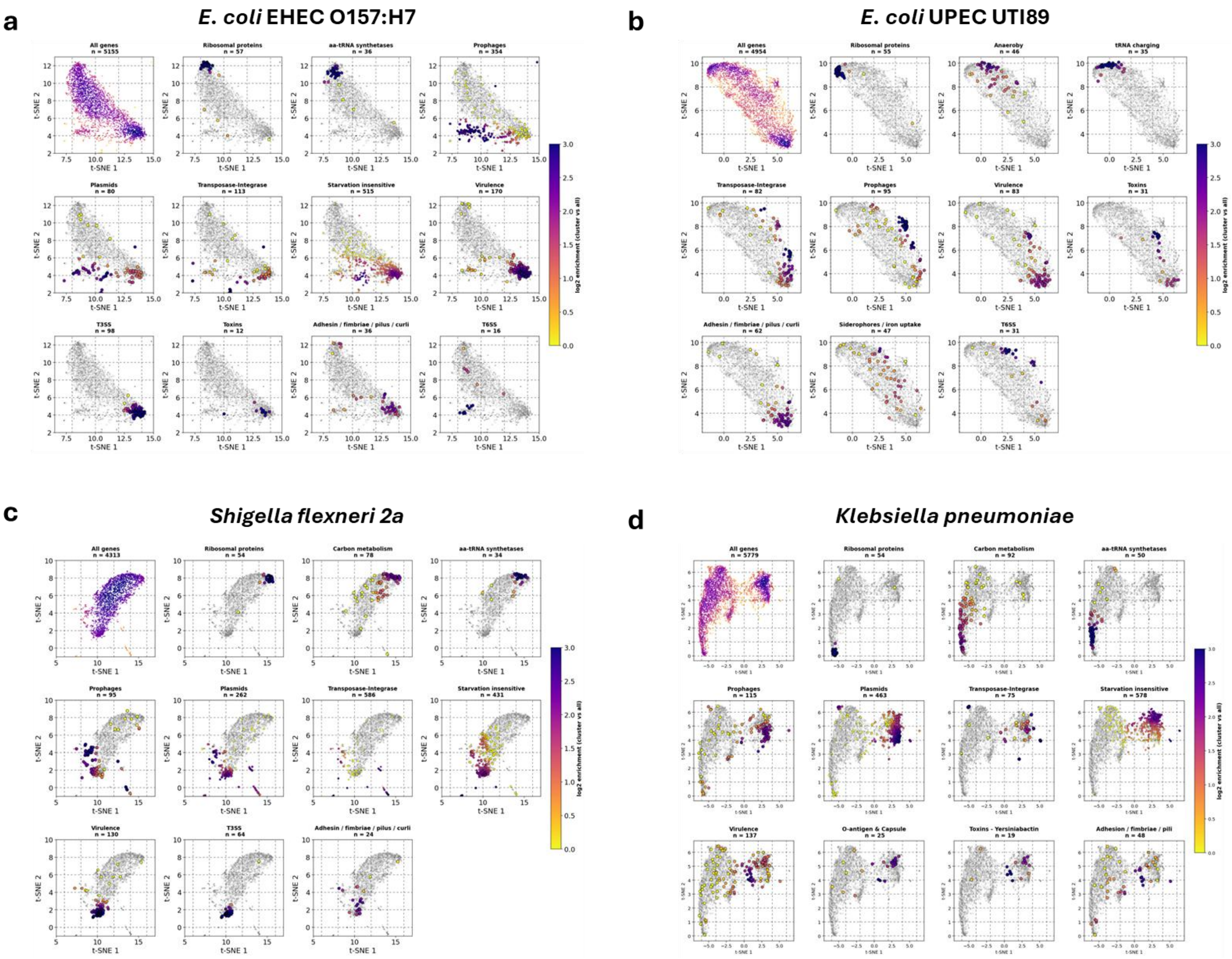
a
E. coli EHEC O157:H7
All genes n = 5155
Ribosomal proteins n = 57
aa-tRNA synthetases n = 36
Prophages n = 354
Plasmids n = 80
Transposase-Integrase n = 113
Starvation insensitive n = 515
Virulence n = 170
T3SS n = 98
Toxins n = 12
Adhesin / fimbriae / pilus / curli n = 36
T6SS n = 16
b
E. coli UPEC UTI89
All genes n = 4954
Ribosomal proteins n = 55
Anaeroby n = 46
tRNA charging n = 35
Transposase-Integrase n = 82
Prophages n = 95
Virulence n = 83
Toxins n = 31
Adhesin / fimbriae / pilus / curli n = 62
Siderophores / iron uptake n = 47
T6SS n = 31
c
Shigella flexneri 2a
All genes n = 4313
Ribosomal proteins n = 54
Carbon metabolism n = 78
aa-tRNA synthetases n = 34
Prophages n = 95
Plasmids n = 262
Transposase-Integrase n = 586
Starvation insensitive n = 431
Virulence n = 130
T3SS n = 64
Adhesin / fimbriae / pilus / curli n = 24
d
Klebsiella pneumoniae
All genes n = 5779
Ribosomal proteins n = 54
Carbon metabolism n = 92
aa-tRNA synthetases n = 50
Prophages n = 115
Plasmids n = 463
Transposase-Integrase n = 75
Starvation insensitive n = 578
Virulence n = 137
O-antigen & Capsule n = 25
Toxins - Yersiniabactin n = 19
Adhesion / fimbriae / pili n = 48
t-SNE 1
t-SNE 2
log2 enrichment (cluster vs all)

**Figure S10. Functional gene clusters generate distinct codon-usage distributions in diverse Enterobacteriaceae pathogens.** Two-dimensional codon-usage UMAP projections for *Escherichia coli* EHEC O157:H7, *E. coli* UPEC UTI89, *Shigella flexneri* 2a and *Klebsiella pneumoniae*. For each pathogen, all genes are shown in grey, together with density overlays for representative functional gene clusters. Density coloring indicates local enrichment relative to the genome-wide gene distribution shown in the first plot of each panel. Functional clusters occupied distinct regions of codon-usage space, supporting the presence of structured and functionally organized codon-usage landscapes across diverse Enterobacteriaceae.

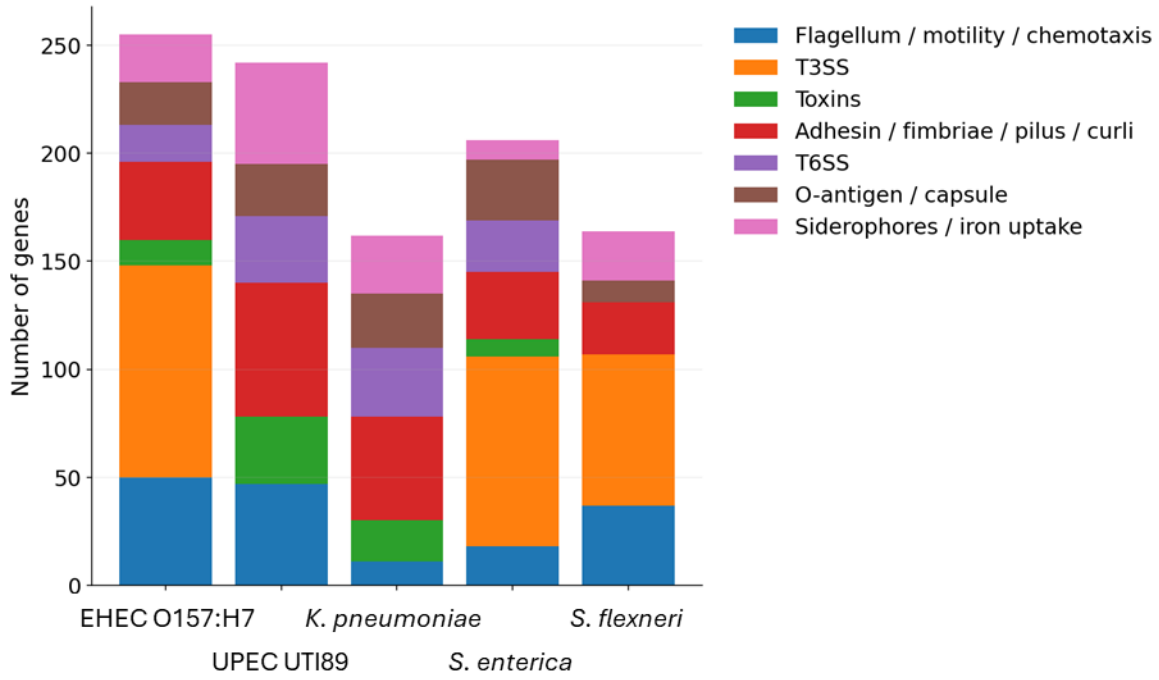


**Figure S11. Selected pathogens cover markedly different virulence strategies.** Virulence-associated genes were identified using ABRicate against curated virulence-factor databases and, for *Klebsiella pneumoniae*, complemented with Kleborate. Genes were manually curated and assigned to functional virulence categories. Gene counts per category were computed for five pathogenic Enterobacteriaceae and plotted as stacked bar plots.

***Escherichia coli* EHEC O157-H7**

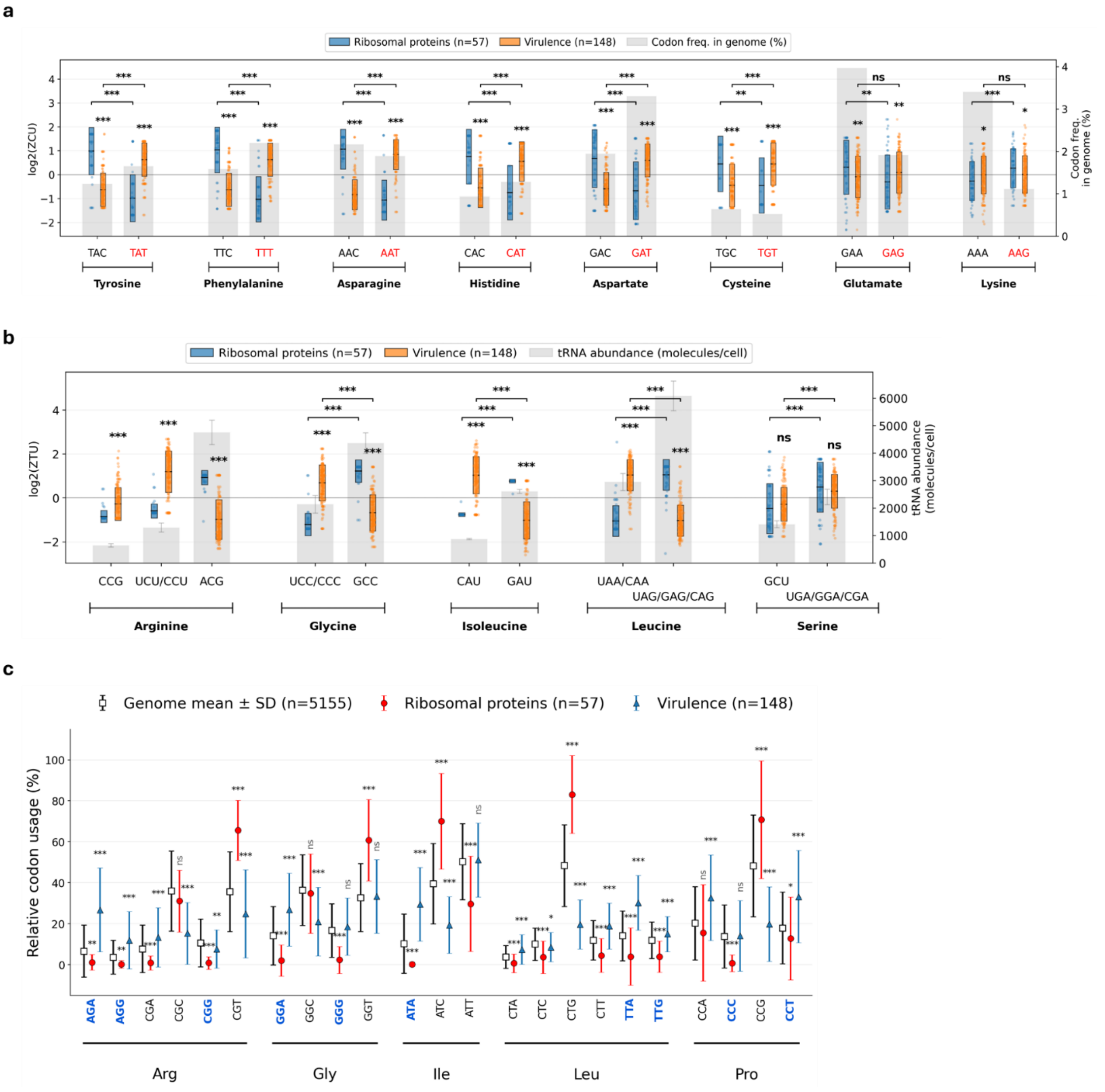

***Escherichia coli UPEC UTI89***

**d**

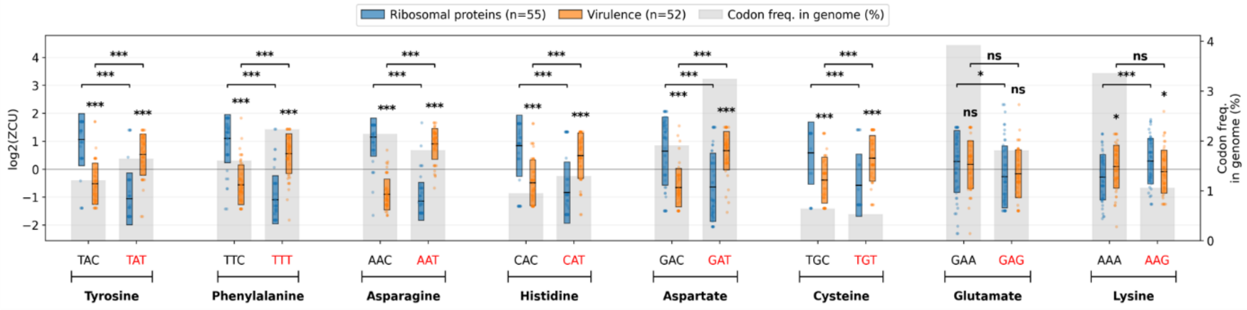


**e**

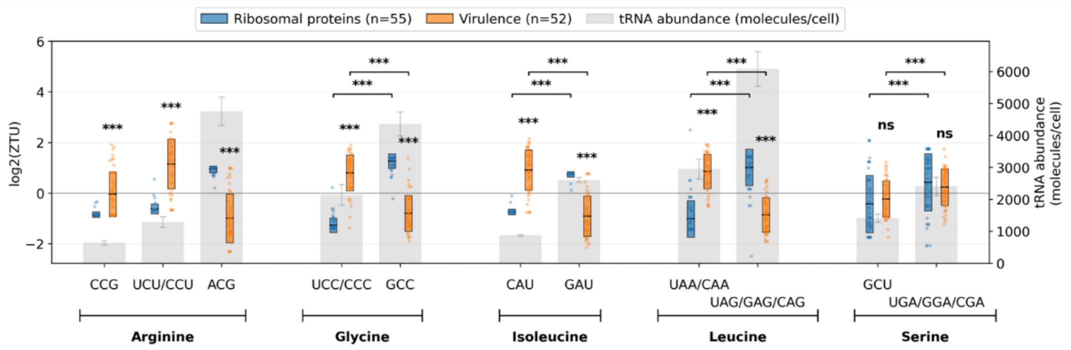


**f**

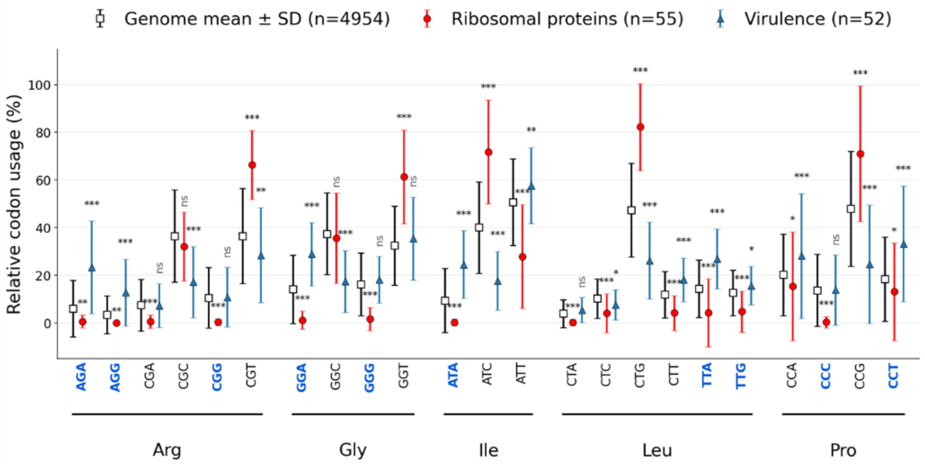

***Shigella flexneri* 2a**

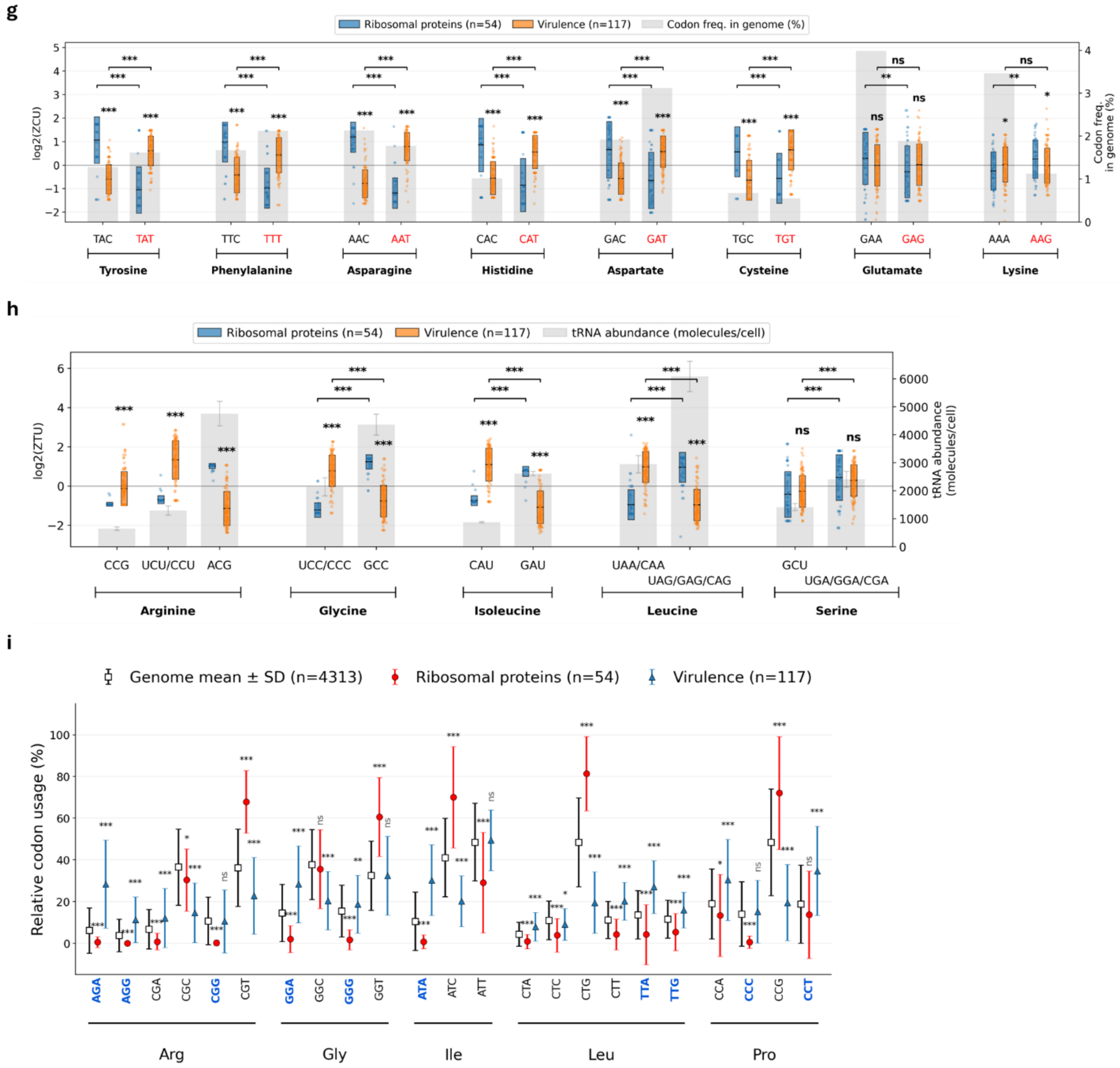

***Klebsiella pneumoniae***

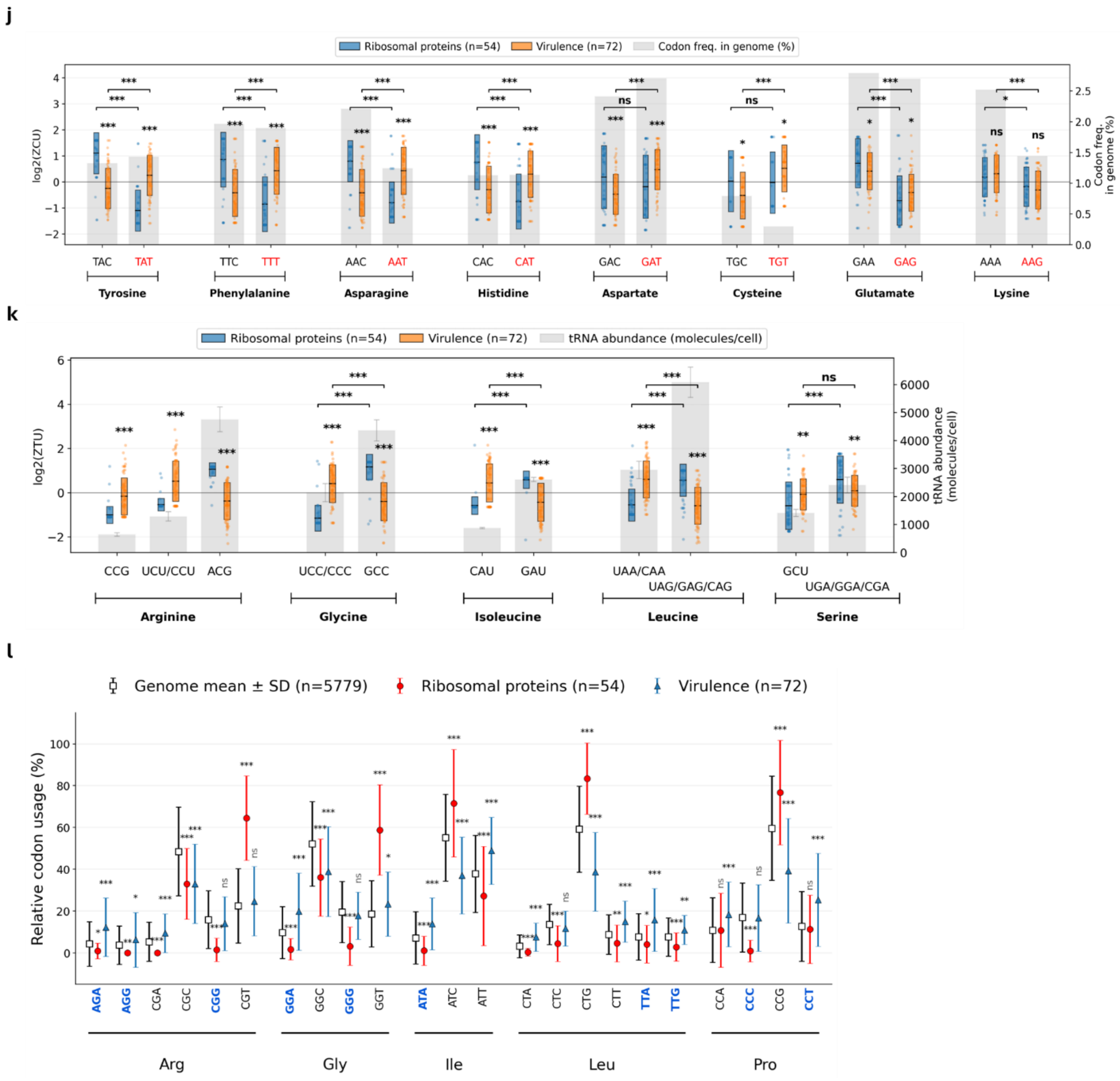


**Figure S12. Virulence genes avoid direct competition with ribosomal genes and favor starvation-insensitive codons in diverse Enterobacteriaceae relevant pathogens.** Data are shown for representative Enterobacteriaceae pathogens including EHEC O157:H7 (a-c), UPEC UTI89 (d-f), *Shigella flexneri* 2a (g-i) and *Klebsiella pneumoniae (j-l)***.** Boxplots show synonymous codon usage z-scores (ZCU) (a, d, g, j) and tRNA usage z-scores (ZTU) (b, e, h, k) for ribosomal and virulence protein genes. (c, f, i, l) Data are average values of ZCU and error bars are SD. Blue codons are starvation insensitive codons, as defined by (1,2). Statistical significance was assessed using two-tailed Student's *t*-tests: * $P < 0.05$, ** $P < 0.01$, *** $P < 0.001$. Note that these data are summarized in Figure 6e.

### Supplementary Tables

| Strains | GC % | Genome RefSeq ID | Family | Gram | Class | Phylum |
|---|---|---|---|---|---|---|
| Acinetobacter baumannii ATCC19606 | 40 | GCF_009035845.1 | Moraxellaceae | – | Gammaproteobacteria | Proteobacteria |
| Agrobacterium tumefaciens str. G3/79 | 59.2 | GCF_013318015.2 | Rhizobiaceae | – | Alphaproteobacteria | Proteobacteria |
| Bacillus subtilis subsp. subtilis str. 168 | 44.2 | GCF_000009045.1 | Bacillaceae | + | Bacilli | Firmicutes |
| Bacteroides fragilis NCTC 9343 | 44.2 | GCF_000025985.1 | Bacteroidaceae | – | Bacteroidia | Bacteroidetes |
| Bdellovibrio bacteriovorus HD100 | 50.7 | GCF_000196175.1 | Pseudobdellovibrionaceae | – | Bdellovibrionia | Bdellovibrionota |
| Brucella abortus 2308 | 58.3 | GCF_000054005.1 | Brucellaceae | – | Alphaproteobacteria | Proteobacteria |
| Burkholderia pseudomallei str. GTC3P0254T | 68.5 | GCF_030297255.1 | Burkholderiaceae | – | Betaproteobacteria | Proteobacteria |
| Caulobacter vibrioides NA1000 (C. crescentus) | 67.7 | GCF_000022005.1 | Caulobacteraceae | – | Alphaproteobacteria | Proteobacteria |
| Chryseobacterium indologenes str. NCTC10796 | 38.3 | GCF_900460995.1 | Weeksellaceae | – | Flavobacteriia | Bacteroidetes |
| Enterobacter cloacae 1382 | 56 | GCF_905331265.2 | Enterobacteriaceae | – | Gammaproteobacteria | Proteobacteria |
| Escherichia coli EHEC O157:H7 str. Sakai | 51.7 | GCF_000008865.2 | Enterobacteriaceae | – | Gammaproteobacteria | Proteobacteria |
| Escherichia coli UPEC UTI89 | 51.6 | GCF_000013265.1 | Enterobacteriaceae | – | Gammaproteobacteria | Proteobacteria |
| Francisella tularensis subsp. novicida D9876 | 33.1 | GCF_000833355.1 | Francisellaceae | – | Gammaproteobacteria | Proteobacteria |
| Helicobacter pylori str. CHC155 | 39.2 | GCF_025998455.1 | Helicobacteraceae | – | Epsilonproteobacteria | Campylobacterota |
| Klebsiella pneumoniae subsp. pneumoniae HS11286 | 58.3 | GCF_000240185.1 | Enterobacteriaceae | – | Gammaproteobacteria | Proteobacteria |
| Listeria monocytogenes EGD-e | 38.4 | GCF_000196035.1 | Listeriaceae | + | Bacilli | Firmicutes |
| Mycobacterium tuberculosis H37Rv | 65.9 | GCF_000195955.2 | Mycobacteriaceae | n/a | Actinomycetes | Actinobacteria |
| Neisseria gonorrhoeae TUM19854 | 53.8 | GCF_013030075.1 | Neisseriaceae | – | Betaproteobacteria | Proteobacteria |
| Neisseria meningitidis Str. PartJ RM8376 | 53.3 | GCF_022869645.1 | Neisseriaceae | – | Betaproteobacteria | Proteobacteria |
| Pseudomonas aeruginosa PAO1 | 67.2 | GCF_000006765.1 | Pseudomonadaceae | – | Gammaproteobacteria | Proteobacteria |
| Pseudomonas putida str. NBRC14164 | 63 | GCF_000412675.1 | Pseudomonadaceae | – | Gammaproteobacteria | Proteobacteria |
| Saccharomyces cerevisiae S288C | 39.6 | GCF_000146045.2 | Saccharomycetaceae | n/a | Saccharomycetes | Ascomycota |
| Salmonella enterica serovar Typhimurium str. SL1344 | 53.4 | GCF_000210855.2 | Enterobacteriaceae | – | Gammaproteobacteria | Proteobacteria |
| Shigella flexneri 2a str. 301 | 51.7 | GCF_000006925.2 | Enterobacteriaceae | – | Gammaproteobacteria | Proteobacteria |
| Staphylococcus aureus subsp. aureus NCTC 8325 | 33.6 | GCF_000013425.1 | Staphylococcaceae | + | Bacilli | Firmicutes |
| Streptococcus pneumoniae str. NCTC7465 | 40.6 | GCF_001457635.1 | Streptococcaceae | + | Bacilli | Firmicutes |
| Streptococcus pyogenes NCTC12064 | 40.4 | GCF_900475035.1 | Streptococcaceae | + | Bacilli | Firmicutes |
| Vibrio cholerae strain RFB16 | 48.2 | GCF_008369605.1 | Vibrionaceae | – | Gammaproteobacteria | Proteobacteria |
| Xanthomonas campestris pv. Raphani str. MAFF106181 | 65.7 | GCF_013388375.1 | Xanthomonadaceae | – | Gammaproteobacteria | Proteobacteria |

**Table S1. List of genomes used in this work, including RefSeq IDs, GC content and phylogenetic information**

| | Ribosomal proteins | tRNA charging | Carbon metabolism | Virulence | SPI-1 | SPI-2 | O-antigen | pSLT | pCol1B9 | Prophages | Transposases Integrases | Starvation insensitive |
|---|---|---|---|---|---|---|---|---|---|---|---|---|
| **Ribosomal proteins** | 1 | 0.00059 | 0.00059 | 0.00059 | 0.00059 | 0.00059 | 0.00059 | 0.00059 | 0.00059 | 0.00059 | 0.00059 | 0.00059 |
| **tRNA charging** | 0.00059 | 1 | 0.00116 | 0.00059 | 0.00059 | 0.00059 | 0.00059 | 0.00059 | 0.00059 | 0.00059 | 0.00059 | 0.00059 |
| **Carbon metabolism** | 0.00059 | 0.00116 | 1 | 0.00059 | 0.00059 | 0.00059 | 0.00059 | 0.00059 | 0.00059 | 0.00059 | 0.00059 | 0.00059 |
| **Virulence** | 0.00059 | 0.00059 | 0.00059 | 1 | 0.28518 | 0.25305 | 0.00615 | 0.00059 | 0.00059 | 0.00059 | 0.00059 | 0.00059 |
| **SPI-1** | 0.00059 | 0.00059 | 0.00059 | 0.28518 | 1 | 0.11152 | 0.01264 | 0.00059 | 0.00059 | 0.00059 | 0.00059 | 0.00059 |
| **SPI-2** | 0.00059 | 0.00059 | 0.00059 | 0.25305 | 0.11152 | 1 | 0.46227 | 0.00059 | 0.00059 | 0.00059 | 0.00059 | 0.00059 |
| **O-antigen** | 0.00059 | 0.00059 | 0.00059 | 0.00615 | 0.01264 | 0.46227 | 1 | 0.00059 | 0.00059 | 0.00059 | 0.00059 | 0.00059 |
| **pSLT** | 0.00059 | 0.00059 | 0.00059 | 0.00059 | 0.00059 | 0.00059 | 0.00059 | 1 | 0.00059 | 0.00059 | 0.00171 | 0.00059 |
| **pCol1B9** | 0.00059 | 0.00059 | 0.00059 | 0.00059 | 0.00059 | 0.00059 | 0.00059 | 0.00059 | 1 | 0.00059 | 0.03947 | 0.00059 |
| **Prophages** | 0.00059 | 0.00059 | 0.00059 | 0.00059 | 0.00059 | 0.00059 | 0.00059 | 0.00059 | 0.00059 | 1 | 0.07927 | 0.00059 |
| **Transposase-Integrase** | 0.00059 | 0.00059 | 0.00059 | 0.00059 | 0.00059 | 0.00059 | 0.00059 | 0.00171 | 0.03947 | 0.07927 | 1 | 0.00059 |
| **Starvation insensitive** | 0.00059 | 0.00059 | 0.00059 | 0.00059 | 0.00059 | 0.00059 | 0.00059 | 0.00059 | 0.00059 | 0.00059 | 0.00059 | 1 |

**Table S2. Matrix of BH-adjusted p-values (p_adj_BH) for all cluster-pair comparisons (diagonal = 1)**

| Component | MW | stock conc. | STOCK VOLUME | | | final conc. | Dilution Factor | Volume: 1L | |
|---|---|---|---|---|---|---|---|---|---|
| MES (pH 3.0) | 195.24 | 1 M | 50ml | 9.76 | g | 100 mM | 10 | 100 | ml |
| KCl | 74.55 | 1 M | 50ml | 3.73 | g | 5 mM | 200 | 5 | ml |
| $NH_4Cl$ | 53.49 | 1.5 M | 50ml | 4.01 | g | 15 mM | 100 | 10 | ml |
| $K_2SO_4$ | 174.26 | 500 mM | 50ml | 4.36 | g | 500 µM | 1000 | 1 | ml |
| $KH_2PO_4$ | 136.09 | 113 mM | 50ml | 0.769 | g | 113 µM | 1000 | 1 | ml |
| $MgSO_4$ | 120.37 | 100 mM | 50 ml | 0.60 | g | 50 µM | 2000 | 500 | µL |
| | 246.48 ($MgSO_4 x 7H_2O$) | | | 1.23 | | | | | |
| Glucose | 180.16 | 20% | 50ml | 10 | g | 0.01% | 2000 | 0.5 | ml |
| Glycerol | 92.09 | 10% | 50ml | 5 | g | 0.02% | 500 | 2 | ml |
| Casamino acids | - | 10% | 50ml | 5 | g | 0 %<br>0.2 % | /<br>50 | 0<br>20 | ml |
| Adjust to pH 5.5 with KOH | | | | | | | | 3 | ml |
| ddH2O | | | | | | | | To 1l | |

**Table S3. Preparation and composition of SPI-2 inducing medium MES-ch.**

| Strain/plasmid | Description | Source |
|---|---|---|
| *Salmonella enterica* serovar Typhimurium SL1344 | WT strain, auxotrophic for histidine | (2) |
| *Salmonella enterica* serovar Typhimurium Sme51 | Prototrophic hisG$^{Leu69}$ derivative of SL1344 | (3) |
| *FreGo911* | *pSC101 kanR PproD-gfpmut2(HEG) PybaJ-mSci* | This study |
| *FreGo914* | *pSC101 kanR PproD-gfpmut2(vir) PybaJ-mSci* | This study |
| *FreGo2001* | *pSC101 kanR PBAD-mtagBFP2(vir)-LVA PybaJ-mNeonGreen* | This study |
| *FreGo2002* | *pSC101 kanR PBAD-mtagBFP2(HEG)-LVA PybaJ-ypet* | This study |
| *FreGo2003* | *pSC101 kanR pBAD-ypet(vir)-LVA PybaJ-mCh* | This study |
| *FreGo2004* | *pSC101 kanR pBAD-ypet(HEG)-LVA PybaJ-mtagBFP2* | This study |
| *FreGo1206* | *pSC101 PssaG-mCh(vir)-LVA PybaJ-mNeonGreen* | This study |
| *FreGo1207* | *pSC101 PssaG-mCh(HEG)-LVA PybaJ-ypet* | This study |
| *FreGo1212* | *pSC101 PssaG-mtagBFP2(vir)-LVA PybaJ-mNeonGreen* | This study |
| *FreGo1213* | *pSC101 PssaG-mtagBFP2(HEG)-LVA PybaJ-ypet* | This study |
| *FreGo1214* | *pSC101 PssaG-ypet(vir)-LVA PybaJ-mCh* | This study |
| *FreGo1215* | *pSC101 PssaG-ypet(HEG)-LVA PybaJ-mtagBFP2* | This study |

**Table S4. Strains and plasmids.** Recoded fluorescent protein genes coding sequences and entire plasmid DNA sequences are provided in Data S2.

| AA | Codon (5'-3') | tRNAs | Anticodon (5'-3') | tRNA molecules per cell (mean) | tRNA molecules per cell (STD) |
|---|---|---|---|---|---|
| Ala | GCA | Ala1/Ala2 | UGC/GGC | 3867 | 287 |
| Ala | GCC | Ala1/Ala2 | UGC/GGC | 3867 | 287 |
| Ala | GCG | Ala1/Ala2 | UGC/GGC | 3867 | 287 |
| Ala | GCU | Ala1/Ala2 | UGC/GGC | 3867 | 287 |
| Arg | CGG | Arg2 | CCG | 639 | 63 |
| Arg | AGA | Arg3/Arg4 | UCU/CCU | 1287 | 166 |
| Arg | AGG | Arg3/Arg4 | UCU/CCU | 1287 | 166 |
| Arg | CGA | Arg1 | ACG | 4752 | 440 |
| Arg | CGC | Arg1 | ACG | 4752 | 440 |
| Arg | CGU | Arg1 | ACG | 4752 | 440 |
| Asn | AAC | Asn1 | GUU | 1193 | 127 |
| Asn | AAU | Asn1 | GUU | 1193 | 127 |
| Asp | GAC | Asp1 | GUC | 2396 | 346 |
| Asp | GAU | Asp1 | GUC | 2396 | 346 |
| Cys | UGC | Cys1 | GCA | 1587 | 126 |
| Cys | UGU | Cys1 | GCA | 1587 | 126 |
| Gln | CAA | Gln1/Gln2 | UUG/CUG | 1645 | 160 |
| Gln | CAG | Gln1/Gln2 | UUG/CUG | 1645 | 160 |
| Glu | GAA | Glu1 | UUC | 4717 | 411 |
| Glu | GAG | Glu1 | UUC | 4717 | 411 |
| Gly | GGA | Gly2/Gly3 | UCC/CCC | 2137 | 320 |
| Gly | GGC | Gly1 | GCC | 4359 | 378 |
| Gly | GGG | Gly2/Gly3 | UCC/CCC | 2137 | 320 |
| Gly | GGU | Gly1 | GCC | 4359 | 378 |
| His | CAC | His1 | GUG | 639 | 95 |
| His | CAU | His1 | GUG | 639 | 95 |
| Ile | AUA | Ile2 | CAU | 868.5 | 23.5 |
| Ile | AUC | Ile1 | GAU | 2605.5 | 70.5 |
| Ile | AUU | Ile1 | GAU | 2605.5 | 70.5 |
| Leu | UUA | Leu4/Leu5 | UAA/CAA | 2944 | 307 |
| Leu | UUG | Leu4/Leu5 | UAA/CAA | 2944 | 307 |
| Leu | CUA | Leu1/Leu2/Leu3 | UAG/GAG/CAG | 6079 | 537 |
| Leu | CUC | Leu1/Leu2/Leu3 | UAG/GAG/CAG | 6079 | 537 |
| Leu | CUG | Leu1/Leu2/Leu3 | UAG/GAG/CAG | 6079 | 537 |
| Leu | CUU | Leu1/Leu2/Leu3 | UAG/GAG/CAG | 6079 | 537 |
| Lys | AAA | Lys1 | UUU | 1924 | 185 |
| Lys | AAG | Lys1 | UUU | 1924 | 185 |
| Met | AUG | Met1 | CAU | 706 | 96 |
| Phe | UUC | Phe1 | GAA | 1037 | 162 |
| Phe | UUU | Phe1 | GAA | 1037 | 162 |
| Pro | CCA | Pro1/Pro2/Pro3 | UGG/CGG/GGG | 2201 | 370 |

| | | | | | |
|---|---|---|---|---|---|
| Pro | CCC | Pro1/Pro2/Pro3 | UGG/CGG/GGG | 2201 | 370 |
| Pro | CCG | Pro1/Pro2/Pro3 | UGG/CGG/GGG | 2201 | 370 |
| Pro | CCU | Pro1/Pro2/Pro3 | UGG/CGG/GGG | 2201 | 370 |
| Ser | AGC | Ser3 | GCU | 1408 | 126 |
| Ser | AGU | Ser3 | GCU | 1408 | 126 |
| Ser | UCA | Ser1/Ser2/Ser4 | UGA/GGA/CGA | 2404 | 283 |
| Ser | UCC | Ser1/Ser2/Ser4 | UGA/GGA/CGA | 2404 | 283 |
| Ser | UCG | Ser1/Ser2/Ser4 | UGA/GGA/CGA | 2404 | 283 |
| Ser | UCU | Ser1/Ser2/Ser4 | UGA/GGA/CGA | 2404 | 283 |
| Thr | ACA | Thr1/Thr2/Thr3/Thr4 | UGU/GGU/CGU | 2656 | 254 |
| Thr | ACC | Thr1/Thr2/Thr3/Thr4 | UGU/GGU/CGU | 2656 | 254 |
| Thr | ACG | Thr1/Thr2/Thr3/Thr4 | UGU/GGU/CGU | 2656 | 254 |
| Thr | ACU | Thr1/Thr2/Thr3/Thr4 | UGU/GGU/CGU | 2656 | 254 |
| Trp | UGG | Trp1 | CCA | 943 | 162 |
| Tyr | UAC | Tyr1 | GUA | 1712 | 257 |
| Tyr | UAU | Tyr1 | GUA | 1712 | 257 |
| Val | GUA | Val1/Val2 | UAC/GAC | 5105 | 411 |
| Val | GUC | Val1/Val2 | UAC/GAC | 5105 | 411 |
| Val | GUG | Val1/Val2 | UAC/GAC | 5105 | 411 |
| Val | GUU | Val1/Val2 | UAC/GAC | 5105 | 411 |

**Table S5. Decoding table for *Escherichia coli*.** This table associates codons with anticodons, tRNA isoacceptors and tRNA isoacceptor abundances extracted from (4).
